# *Deinococcus radiodurans* HU: a versatile architect of nucleoid structure and plasticity

**DOI:** 10.64898/2026.08.18.745388

**Authors:** Harald Bernhard, Salvatore De Bonis, Anne-Sophie Banneville, Maria Bacia-Verloop, Lorenzo Gaifas, Brandon Le Bon, Georg Wolff, Michael Hall, Roy González-Alemán, François Dehez, Irina Gutsche, Joanna Timmins

## Abstract

Nucleoid-associated proteins are known to compact and organise bacterial genomes, but how their local DNA binding shapes higher-order chromatin architecture remains unclear. Using *in vitro* and *in situ* approaches, we investigated how *Deinococcus radiodurans* HU (*Dr*HU) orchestrates nucleoid organisation. *Dr*HU engages DNA through two synergistic binding modes: high-affinity β-hairpin-mediated stabilisation of DNA loops and low-affinity, multivalent interaction via its N-terminal tail, driving parallel, uniformly spaced DNA alignment. *In vitro, Dr*HU organises supercoiled plasmid DNA into regular 2D lattices of double-spirals, while *in situ*, UV-C light induces a nucleoid transition from a loose DNA mesh to a blue liquid crystalline phase characterised by DNA swirls. The striking structural similarity and shared inter-filament spacing of these two arrangements suggests that *Dr*HU plays a pivotal role in genome organisation through its versatile binding properties that enable DNA loop stabilisation, protection and DNA bridging, linking molecular interactions to 3D genome ordering and stress-induced remodelling.

## Introduction

Understanding how cells compact and organise their genetic material while preserving efficient and regulated access to genetic information remains a central question in biology. While eukaryotic chromatin organisation has been characterised in considerable detail, the bacterial nucleoid, despite its apparent simplicity, remains far less well understood. This disparity stems from fundamental differences in genome organisation. In eukaryotes, nucleosomes provide a well-defined building block of chromatin architecture^1–5^ that facilitates its characterisation across scales through biochemical, structural, imaging and genome-wide approaches. In contrast, the bacterial nucleoid lacks such a defined unit, contains roughly tenfold less protein per DNA mass^6,7^, and relies instead on nucleoid-associated proteins (NAPs), DNA supercoiling and macromolecular crowding for compaction^8^.

NAPs include a diverse group of small, basic architectural proteins including HU, Fis, IHF and H-NS that bind, bend and/or bridge DNA^9^. Among these, HU is the most abundant and widespread. It assembles as either homo- or heterodimers^10,11^, and has been shown to bind strongly to distorted DNA regions through conserved β-hairpin arms and more weakly along unconstrained DNA via three conserved lysine residues^10,12^. These distinct binding modes suggest that HU may have multiple functions in DNA organisation^13^. However, despite decades of biochemical and structural studies, the mechanisms of HU-mediated nucleoid organisation remain difficult to rationalise because of variations in HU isoforms and in the number and redundancy of NAPs across species, leaving open how different modes of local HU-DNA binding translate into larger-scale genome organisation.

*Deinococcus radiodurans* is a non-pathogenic gram-positive bacterium well-known for its extraordinary resistance to DNA damaging agents^14,15^. Under normal growth conditions, *D. radiodurans* nucleoids are highly dynamic, capable of adopting various morphologies as a function of the cell cycle^16^. In response to intense UV-C light, *D. radiodurans* nucleoids transition to a compact, spherical morphology within 1 hour, before decompacting as cells recover^17^. Interestingly, in *D. radiodurans,* HU is essential for viability and constitutes the unique small architectural NAP^18–20^; it is therefore likely to be a key player in *D. radiodurans* nucleoid organisation. Depletion experiments have indeed shown that reduced levels of *D. radiodurans* HU (*Dr*HU) rapidly lead to nucleoid decompaction^20^. This is further supported by recent single-molecule tracking data demonstrating a marked change in *Dr*HU dynamics in UV-irradiated cells, which closely correlates with the observed changes in nucleoid morphology^17^. Unlike many of its homologues, *Dr*HU possesses an unusual lysine-rich N-terminal tail (N-tail), reminiscent of linker histone H1 that plays a critical role in DNA compaction in eukaryotes^21^, which may further modulate DNA binding, bending and bridging properties of *Dr*HU.

Here, we combine cryo-electron microscopy (cryo-EM), biochemical and mutational studies, molecular dynamics (MD) simulations and *in situ* cryo-electron tomography (cryo-ET), to elucidate the versatile DNA-binding modes of *Dr*HU and how they drive higher-order nucleoid organisation and stress-induced genome rearrangements in *D. radiodurans*. This work provides a mechanistic and structural framework for a broader understanding of the key roles of HU proteins in bacterial nucleoid structure and plasticity.

## Results

### *Dr*HU organises supercoiled DNA into a regular, spiral arrangement

To gain insight into how *Dr*HU binds DNA and organises the genome, we mixed supercoiled (SC-pUC19) or linear (L-pUC19) pUC19 plasmid DNA, used as a genomic DNA mimic, with *Dr*HU at different protein:DNA ratios, and imaged the mixtures by cryo-EM. Electrophoretic mobility shift assays (EMSA) confirmed similar binding profiles to both plasmid topologies (Supplementary Fig. S1). At a 100:1 (*Dr*HU:SC-pUC19) ratio, supercoiled DNA adopted a quasi-regular 2D spiral-like arrangement with a uniform spacing of ∼9 nm between adjacent DNA filaments (Fig. 1a, d). The apparent planarity of these assemblies likely results from embedding in a very thin layer of vitrified ice and interaction with the air-water interface. At lower ratios, only shapes resembling SC-pUC19 alone were observed, occasionally decorated with small dark densities (Supplementary Fig. S2a-Fig. 3a). At higher ratios, sample precipitation hampered conclusive evaluation. Although the spiral pattern was not observed when *Dr*HU was mixed with L-pUC19 (Fig. 1b and Supplementary Fig. S2b), the linear DNA adopted a wavy, locally parallel arrangement with a similar ∼9 nm inter-filament spacing, as observed for SC-pUC19 spirals (Fig. 1b, d). Importantly, spiral or parallel DNA arrangements were never observed in pUC19 samples in the absence of *Dr*HU (Supplementary Fig. S3a-b).

**Figure 1:**
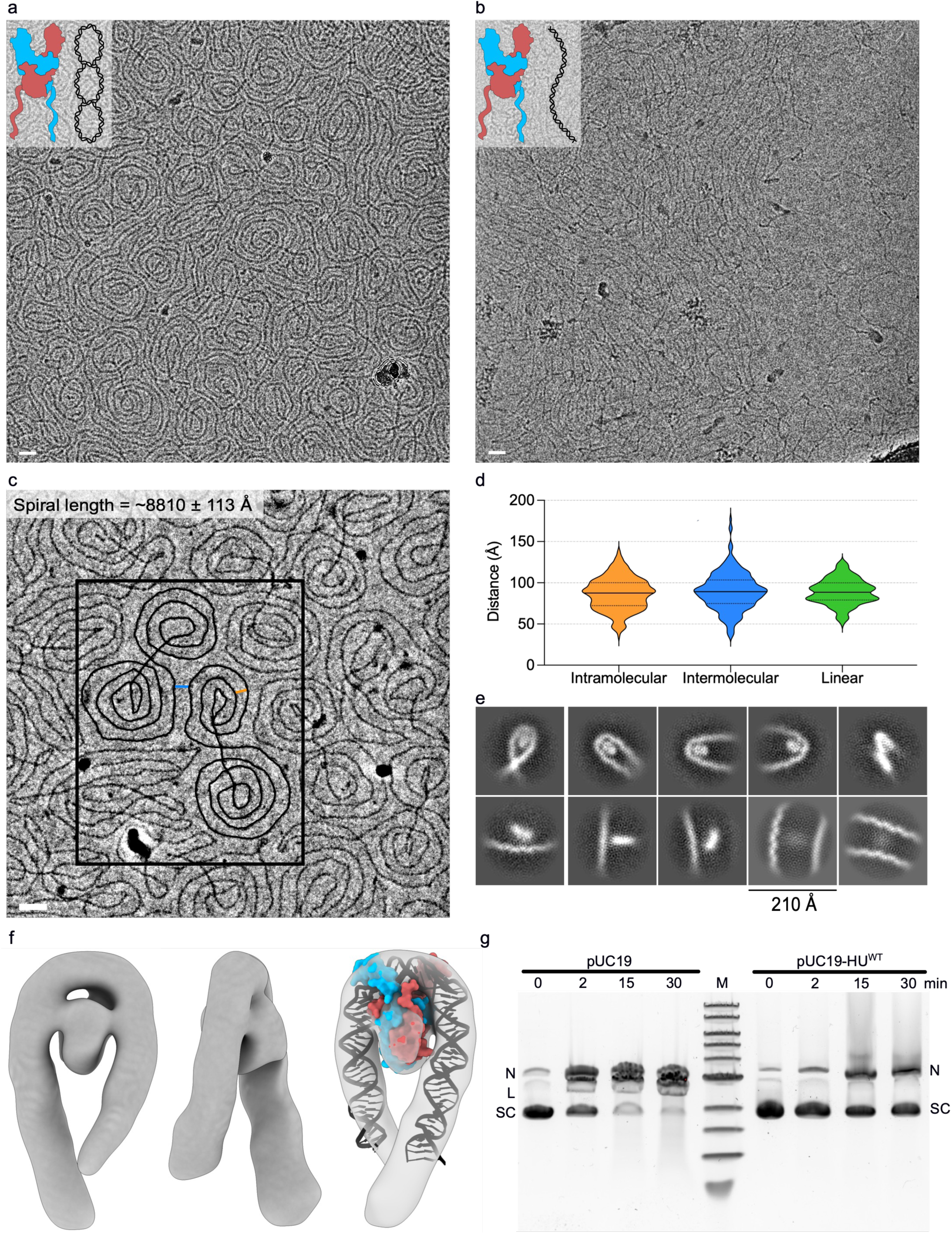
*Dr*HU induces higher-order 2D arrangement of linear and supercoiled plasmid DNA. **(a)-(b)** Representative cryo-EM micrographs of SC-pUC19 **(a)** and L-pUC19 **(b)** together with *Dr*HU. Scale bar: 20 nm. **(c)** Representative cryo-EM micrograph from the phase plate data collection with highlighted sc-pUC19 plasmid (black), inter-filament spacing of DNA within a given plasmid (orange) and between adjacent plasmids (blue). **(d)** Violin plot representing inter-filament distance measurements of intramolecular (orange; N=267) and intermolecular (blue; N=231) SC-pUC19 as well as L-pUC19-HU (green; N=140). **(e)** Examples of 2D class averages of *Dr*HU within DNA loops or bound more loosely to straight or curved DNA showing more diffuse signal. **(f)** Low resolution 3D map of the *Dr*HU dimer (red & blue) bound within a DNA loop (black). An AlphaFold3 model of a *Dr*HU dimer bound to DNA was fitted into the EM map. **(g)** Agarose gel electrophoresis of the products of the S1 nuclease assay, alongside nicked (N), linear (L) and supercoiled (SC) pUC19 plasmid DNA, as indicated. SC-pUC19 alone (left) or in the presence of *Dr*HU (right) was treated with S1 nuclease for 0, 2, 15 or 30 min.

Intrigued by the remarkable DNA spirals formed by *Dr*HU:SC-pUC19 at a 100:1 ratio, we acquired a cryo-EM dataset using a phase plate to enhance contrast and improve the likelihood of identifying *Dr*HU (Fig. 1c). DNA spirals were found to frequently adopt an eight-shaped double spiral arrangement (Fig. 1a, c) with an average length of 8810 ± 11 Å (N=21), close to the theoretical contour length of a pUC19 plasmid (9130 Å), suggesting that each of these structures corresponds to an individual pUC19 molecule. The mean inter-filament distance within a single plasmid and between adjacent plasmids was measured to be 87 ± 20 Å (N=267) and 90 ± 23 Å (N=231) respectively, indicating a uniform inter-filament spacing within and between plasmid molecules (Fig. 1d). Similarly, the distance between parallel *Dr*HU:L-pUC19 DNA filaments was found to be in the same range (89 ± 15 Å; N=140) (Fig. 1d). These results strongly suggest that the regular DNA spacing is caused by *Dr*HU, acting as a distance holder between neighbouring DNA filaments.

### *Dr*HU exhibits multiple DNA binding modes and stabilises tight DNA loops

2D class averaging of 31,949 DNA segments bearing putative *Dr*HU densities revealed several different DNA binding modes (Fig. 1e). Well-defined *Dr*HU densities were identified within tight DNA loops, while more diffuse densities were instead positioned along straight or slightly curved DNA duplexes, at variable angles relative to the DNA axis, suggestive of a more dynamic binding mode or partial denaturation at the air-water interface (Fig. 1e). A few 2D classes also showed weak inter-filament density, indicating a potential DNA bridging mode (Fig. 1e). Only loop-containing classes corresponded to views of DNA-bound *Dr*HU from multiple orientations, suitable for 3D reconstruction (Supplementary S 4). However, tight loops were rare (typically 2 to 4 per plasmid; Supplementary Fig. S5a-c) thereby limiting the total number of loop-containing particles (11,324) and restricting the achievable resolution, further constrained by the low molecular weight of a *Dr*HU dimer (24.6 kDa). Nevertheless, an AlphaFold3 model of the core domain of a *Dr*HU homodimer bound to a strongly bent DNA and lacking the flexible N-tails of *Dr*HU, showed a very good agreement with our experimentally derived cryo-EM map (Fig. 1f).

In supercoiled DNA, the mechanical tension in tight turns is partially relieved by local distortion, unwinding or base pair-disruption, leading to the flipping out of one or more nucleotides and the formation of short single-stranded stretches, susceptible to cleavage by certain nucleases^22,23^. S1 and Bal31 nuclease assays confirmed that *Dr*HU binding to tight DNA turns protects supercoiled DNA from cleavage (Fig. 1g and Supplementary Fig. S5d) and likely further stabilises this constrained conformation. HU proteins have also been reported to bend DNA^24^, raising the possibility that *Dr*HU might not only protect but also actively introduce tight loops. However, the mean number of loops per SC-pUC19 plasmid bound to *Dr*HU in our cryo-EM images (3.1; N=153) matches values derived from AFM observations of naked supercoiled pUC18^25^, making this scenario unlikely.

### β-hairpins, N-tails and tetramerisation of *Dr*HU contribute to DNA binding and bridging

The cryo-EM map of the *Dr*HU-DNA loop indicates that the β-hairpin arms of the *Dr*HU dimer are required for this binding mode in agreement with earlier crystal structures^26–29^, showing that the β-hairpins insert into the minor groove and induce DNA kinks^24^. However, the putative role of the N-tail of *Dr*HU remained to be elucidated. To determine the respective roles of these two regions of *Dr*HU in DNA binding and higher-order genome organisation, we generated three *Dr*HU variants: a β-hairpin deleted form of *Dr*HU (HU^ΔHp^), a N-tail truncated form of *Dr*HU (HU^Δ28^), and a variant missing both of these features (HU^Δ28ΔHp^). We then compared their DNA binding properties to those of wild-type *Dr*HU (HU^WT^). EMSA revealed two *Dr*HU binding modes: high-affinity loop binding (Kd ∼0.1 µM) dependent on the β-hairpin and non-specific, low-affinity DNA binding (Kd ∼2.5 µM) requiring the N-tail (Fig. 2a). Cryo-EM observations confirmed that β-hairpin deletion disrupted tight loop binding and spiral formation, and further demonstrated that the N-tail is needed for uniform DNA spacing (Fig. 2b–c and Supplementary Fig. S6). Accordingly, both modes of DNA binding were affected in the double mutant, although some residual DNA binding was still observed (Fig. 2a, d). Fluorescence polarisation (FP) measurements confirmed that the flexible N-tails of *Dr*HU constitute the primary, albeit lower-affinity, determinant of *Dr*HU binding to DNA, responsible for establishing the uniform inter-filament spacing, whereas the β-hairpins mediate fewer, but stronger interactions localised to tight DNA turns (Fig. 2d).

**Figure 2:**
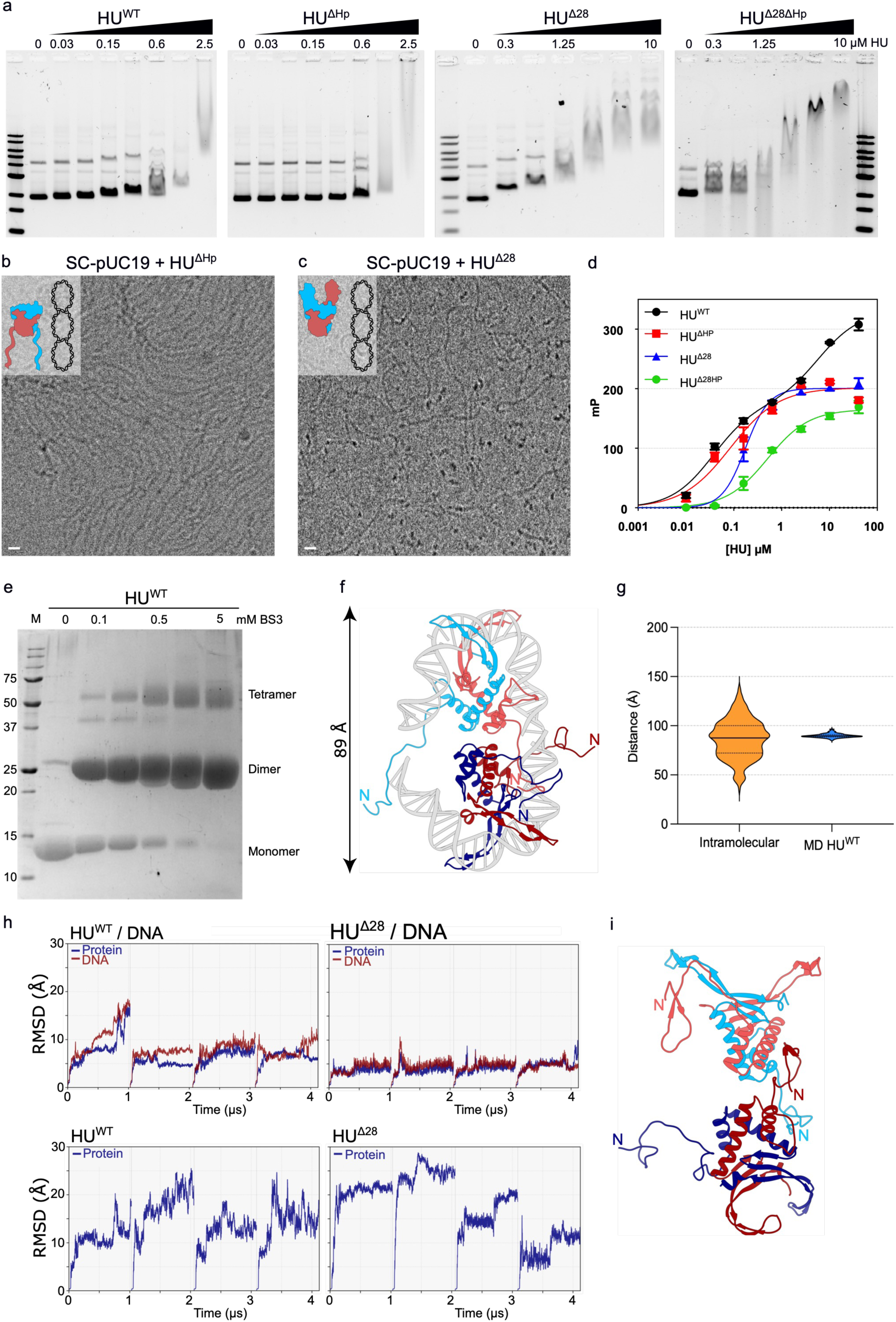
DNA binding properties of wild-type versus mutant *Dr*HU. **(a)** EMSA analysis of 0-2.5 µM (or 10µM for the double mutant) *Dr*HU (HU^WT^, HU^ΔHp^, HU^Δ28^ and HU^Δ28ΔHp^) binding to SC-pUC19 plasmid. **(b)-(c)** Examples of cryo-EM micrographs of HU^ΔHp^ **(b)** and HU^Δ28^ **(c)** variants bound to SC-pUC19 plasmid DNA. **(d)** Fluorescence polarisation titration curves of *Dr*HU binding to dsDNA (50 mer). Data points are the mean of 3 independent measurements and the error bars correspond to the standard deviation. The data points were fitted to a one- or a two-site specific binding model with Hill slope in Graph Pad Prism 10. **(e)** SDS-PAGE analysis of BS3 chemically crosslinked *Dr*HU, revealing the formation of both dimers and tetramers. **(f)** Representative model of tetrameric *Dr*HU bound to dsDNA derived from AlphaFold3 prediction and MD simulations. Each dimer is bound to a 36bp DNA duplex (grey). One dimer is coloured in light blue/red, and the other in dark blue/red. The distance between the β-hairpin structures of each dimer is ∼89 Å. *Note: although this particular DNA configuration is unlikely to occur with longer DNA as in the spiral arrangement, this model provides important insight into the role of the N-tails in DNA binding and tetramer formation.* **(g)** Violin plot comparing inter-filament distance measurements obtained for HU-DNA spirals (orange; N=267) with the DNA-DNA distances derived from MD simulations of *Dr*HU^WT^ (blue) tetramers bound to dsDNA (N=4120). **(h)** RMSD profiles tracking the protein (dark blue lines) and DNA (red lines) dynamics of the four explored systems over a 4.12 μs total trajectory divided into four replicates separated by dashed vertical lines. **(i)** Representative model of tetrameric *Dr*HU^WT^ in its apo state stabilised by contacts mediated by the helical core domains and the N-tails.

In the FP experiments, HU^WT^, unlike the mutant variants, exhibited a two-phase binding profile when incubated with 50 mer dsDNA (Fig. 2d). We hypothesised that the second binding event may reflect the formation of a higher oligomeric state of *Dr*HU on DNA. Chemical cross-linking revealed that, in addition to the expected homodimers, all *Dr*HU variants form tetramers, though to varying extents: ∼20% for HU^WT^, reduced to 10-15% for the single deletion mutants and to 5% in the double mutant (Fig. 2e and Supplementary Fig. S7a-b). These results indicate that the N-tail, and to a lesser extent, the β-hairpin, contribute to tetramer formation. Consistently, MD simulations performed using an AlphaFold3 model of tetrameric *Dr*HU bound to DNA as a starting model (Supplementary Fig. S7c) indicates that a back-to-back tetramer, stabilised through contacts involving the helical core domain and the N-tails of *Dr*HU, could bridge DNA molecules with an inter-DNA distance of ∼89 Å, in close agreement with the experimentally observed inter-filament distance (Fig. 2f-g and Supplementary Fig. S7d). Additionally, the MD simulations reveal a key role of the N-tails in modulating the dynamic behaviour of *Dr*HU (Fig. 2h-i and Supplementary Fig. S8). In the absence of DNA, the N-tails stabilise the tetramer by providing additional contact points between the dimers in addition to the weak contacts between the helical core domains of *Dr*HU dimers (Fig. 2i and Supplementary Fig. S7 and S8), whereas in the presence of DNA, the N-tails significantly increase the conformational landscape of *Dr*HU-DNA assemblies (exemplified by significantly increased RMSD values along the runs) by engaging in transient and multivalent interactions with both the DNA and the neighbouring *Dr*HU dimer (Fig. 2h and Supplementary Fig. S8). The highly dynamic nature of *Dr*HU-DNA assemblies and *Dr*HU tetramers is reflected by the diffuse densities observed in the 2D class averages of *Dr*HU loosely bound to DNA via its N-tail (Fig. 1e).

### Exposure of *D. radiodurans* cells to UV-C light drives the nucleoid into a blue liquid crystalline phase

Finally, we wondered how the structural insights from the *in vitro* assembled *Dr*HU-DNA data can advance our understanding of bacterial nucleoid architecture, and whether the *Dr*HU-driven double spirals of supercoiled DNA and their supramolecular 2D assemblies reflect nucleoid organisation *in situ*. Thus, building on our recent fluorescence microscopy data showing that UV-C irradiation induces compaction of the *D. radiodurans* nucleoid^17^, we performed comparative cryo-ET imaging on ∼180 nm thick cryo-FIB-milled lamella of exponentially growing bacteria, plunge-frozen either before or 1h after exposure to UV-C light (Fig. 3a-b). Cryo-ET of UV-C-irradiated cells showed a majority of nucleoids (∼90%) transitioning from a loosely organised, irregular DNA mesh (Fig. 3a; Supplementary Video 1) to distinctive swirls with locally aligned and uniformly spaced DNA filaments (spacing ∼8.5 ± 1.5 nm, N=200; Fig. 3b-c; Supplementary Video 2), matching the inter-filament spacing measured in the *Dr*HU:SC-pUC19 double spirals (∼9 nm; Fig. 1d). This close agreement between *in situ* and *in vitro* measurements hints at a comparable underlying DNA organisation.

**Figure 3:**
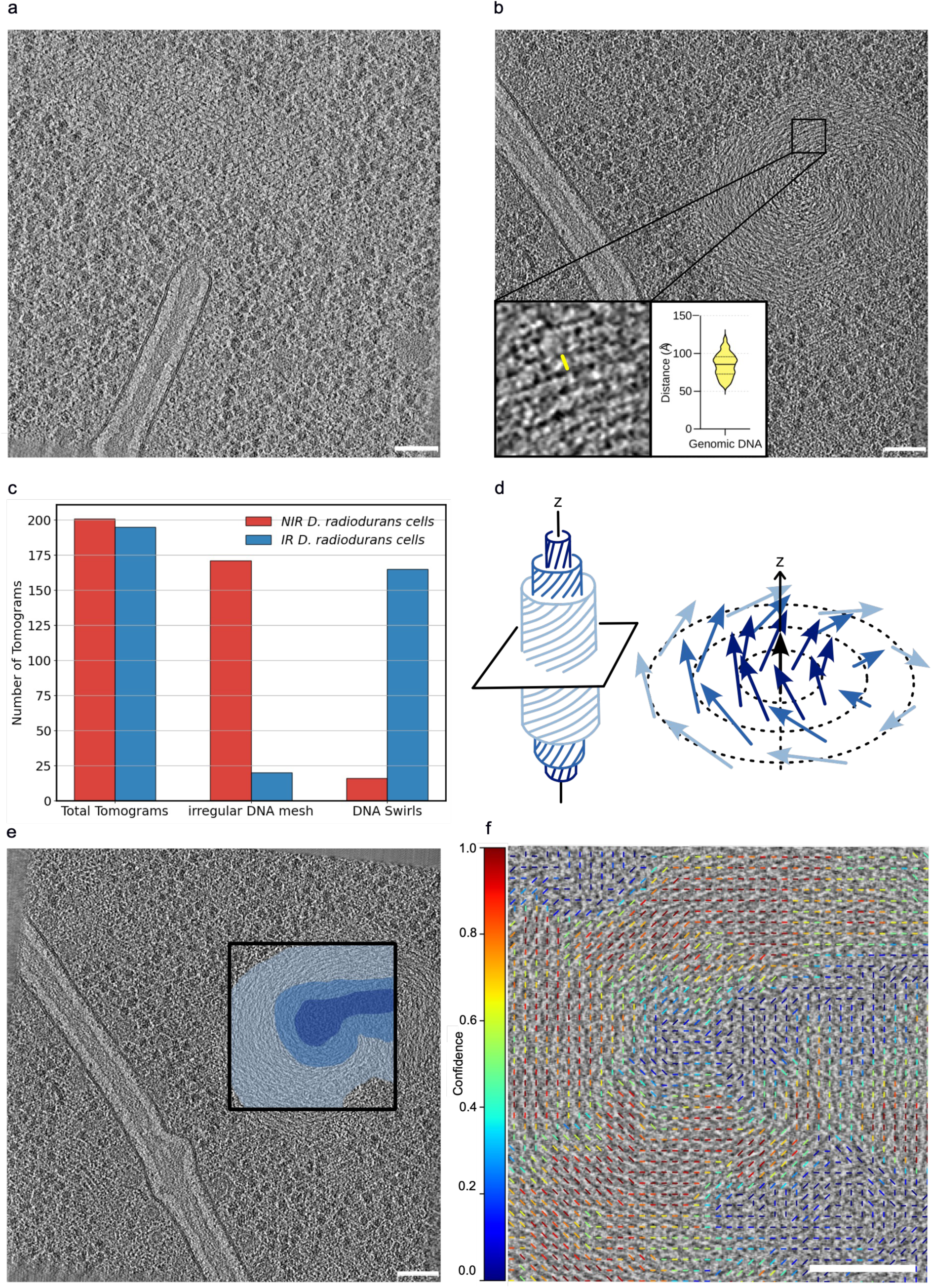
UV-C light induces nucleoid reorganisation into a blue liquid crystalline phase. **(a)-(b)** Slice through a representative tomogram of a non-irradiated **(a)** and UV-irradiated *D. radiodurans* cell **(b)**. **(c)** Histogram plot of the occurrence of DNA swirls versus loose DNA in tomograms of non-irradiated (red) and irradiated (blue) cells. **(d)** Schematic diagram illustrating the organisation of DNA filaments in a double-twist cylinder, a building block of a blue liquid crystalline phase. **(e)** Two examples of UV-irradiated *D. radiodurans* nucleoids bearing DNA swirls with double-twist cylinder features, highlighted in light to dark blue. **(f)** Close-up view of the boxed region in e, overlaid with the confidence scores of the corresponding orientation field, which exhibits a clear spiral arrangement. In regions of the swirl where DNA filaments are readily distinguishable and locally aligned, filament orientation can be reliably estimated, resulting in high confidence scores (red). Conversely, where filaments are less distinct or less ordered, orientation estimates are less reliable and confidence scores decrease (green/blue), notably near the axis of the double-twist cylinder, where filaments are aligned along the cylinder axis while image gradients are computed in the plane perpendicular to it. Scale bars: 75 nm.

Examination of the orientation fields of DNA filaments computed from local image-intensity variations revealed a clear spiralling organisation (Fig. 3e-f), providing an additional line of evidence, alongside tomographic appearance (Fig. 3b) and Fourier analysis (Supplementary Fig. S9), that these DNA swirls are remarkably reminiscent of double-twist cylinders, the fundamental building blocks of blue phases of chiral liquid crystals. In a double-twist cylinder, DNA filaments are aligned parallel to the cylinder axis at the centre, but progressively tilt outward with increasing radial distance, giving rise to a continuous, azimuthally symmetric twist in the cross-section (Fig. 3d-f; Supplementary Fig. S10; Supplementary Video 3)^30^. A blue phase then arises from packing of double-twist cylinders into a 3D lattice^31^. In irradiated *D. radiodurans* cells, one or more DNA swirls were seen in single nucleoids (Fig. 3e; Supplementary Fig. S10). Interestingly, although visually less prominent, comparable DNA patterns were observed in early CEMOVIS (Cryo-EM Of Vitreous Sections) images of stationary-phase or stressed bacteria, in particular in *D. radiodurans*^7,33^, and were interpreted as evidence that bacterial nucleoids adopt a pre-cholesteric or blue phase-like liquid crystalline state^34^ under stress conditions^7,33^. However, these studies reported an inter-DNA spacing of ∼4-6 nm, substantially lower than the ∼8-9 nm spacing observed in our swirls and spirals. This discrepancy may reflect differences in the measurement approaches, since the previous CEMOVIS measurements were performed on 2D projection images of ∼70 nm thick sections and therefore likely affected by the superposition of multiple DNA layers, whereas our measurements were done on 7.872 Å thick tomographic slices, reducing projection artefacts and providing a more direct estimate of the true 3D DNA spacing and organisation.

## Discussion

In this study, we combined cryo-EM, cryo-ET, biochemical assays and MD simulations to dissect the multifunctional roles of *Dr*HU as a DNA binder, bender and bridger, and determine the respective contributions of distinct structural elements, such as N-terminal tail, β-hairpin and tetramerisation, to these different architectural properties.

Previous studies identified two primary DNA binding modes for HU proteins. First, a high-affinity, structure-specific binding mode dependent on β-hairpin insertion into the minor groove, observed in numerous structural studies of HU homologues^26–29^ is likely responsible for the preference of HU for binding to distorted DNA substrates *in vitro*, such as four-way junctions, DNA repair intermediates or gapped DNA^35–38^. In contrast, the second mode of DNA binding is believed to be non-specific and to result from lateral binding of HU along the DNA filament through electrostatic interactions between surface exposed lysine residues (Lys3, Lys18, Lys83 in *E. coli* HUα) on the helical core and the DNA backbone^39–41^. Unlike the tight binding mediated by the β-hairpin structures of dimeric HU, this alternative binding mode has been reported to result in extended nucleoprotein filament formation and transition from dimer to higher order oligomerisation states of HU^40,42,43^. Furthermore, in a recent study, Chan *et al.* used molecular dynamics simulations to show that the β-hairpin arms of *E. coli* HU can also function as antennae for the initial binding to DNA and to facilitate displacement along and between DNA segments, in conjunction with the low-affinity, non-specific binding along the DNA axis, mediated by the helical core^44^.

Here, we report that *Dr*HU interacts with DNA through at least two distinct, yet synergistic, binding modes, each with unique structural determinants and functional consequences (Fig. 4). We identified a first mode involving high-affinity binding of *Dr*HU to tight DNA loops, a process critically dependent on its β-hairpin structure and similar to that reported in several earlier structural studies of HU homologues from various bacterial species^26–29^. It relies on the insertion of the two β-hairpin structures from a single *Dr*HU homodimer into two successive minor grooves of the DNA and is functionally significant. It notably protects supercoiled DNA from nuclease cleavage and likely stabilises this energetically unfavourable conformation of the DNA, thereby alleviating topological constraints associated with DNA supercoiling. Like other HU homologues, *Dr*HU exhibits a second mode of DNA binding, which is much more dynamic and mediated by multivalent electrostatic interactions between surface exposed lysine residues and the DNA backbone and results in a wide diversity of DNA binding configurations, as illustrated by the diverse 2D classes of *Dr*HU-DNA assemblies and the multiple conformations sampled during the MD simulation runs. Whilst *E. coli* HU uses lysine residues within its helical core for this interaction^39,40^, *Dr*HU relies on its disordered, lysine-rich N-terminal tail to fulfil this function. Indeed, deleting the N-tail leads to severely impaired DNA binding, suggesting that this lower-affinity binding mode constitutes the main determinant of *Dr*HU DNA binding. As a result, a vast majority of DNA-bound *Dr*HU is flexibly tethered to the DNA and adopts a range of orientations relative to the DNA axis (parallel, orthogonal or intermediate). It should be noted, however, that although the double mutant missing both the N-tail and the β-hairpin exhibited severely impaired DNA binding, weak DNA binding was nonetheless retained, suggesting that a third region of *Dr*HU, possibly its helical core, may also contribute to non-specific DNA binding. *Dr*HU lacks several surface-exposed lysine residues present in other HU homologues, including the equivalent of *E. coli* K83, which may explain its reliance on the N-tail, and to a lesser extent on its helical core, for non-specific DNA binding.

**Figure 4:**
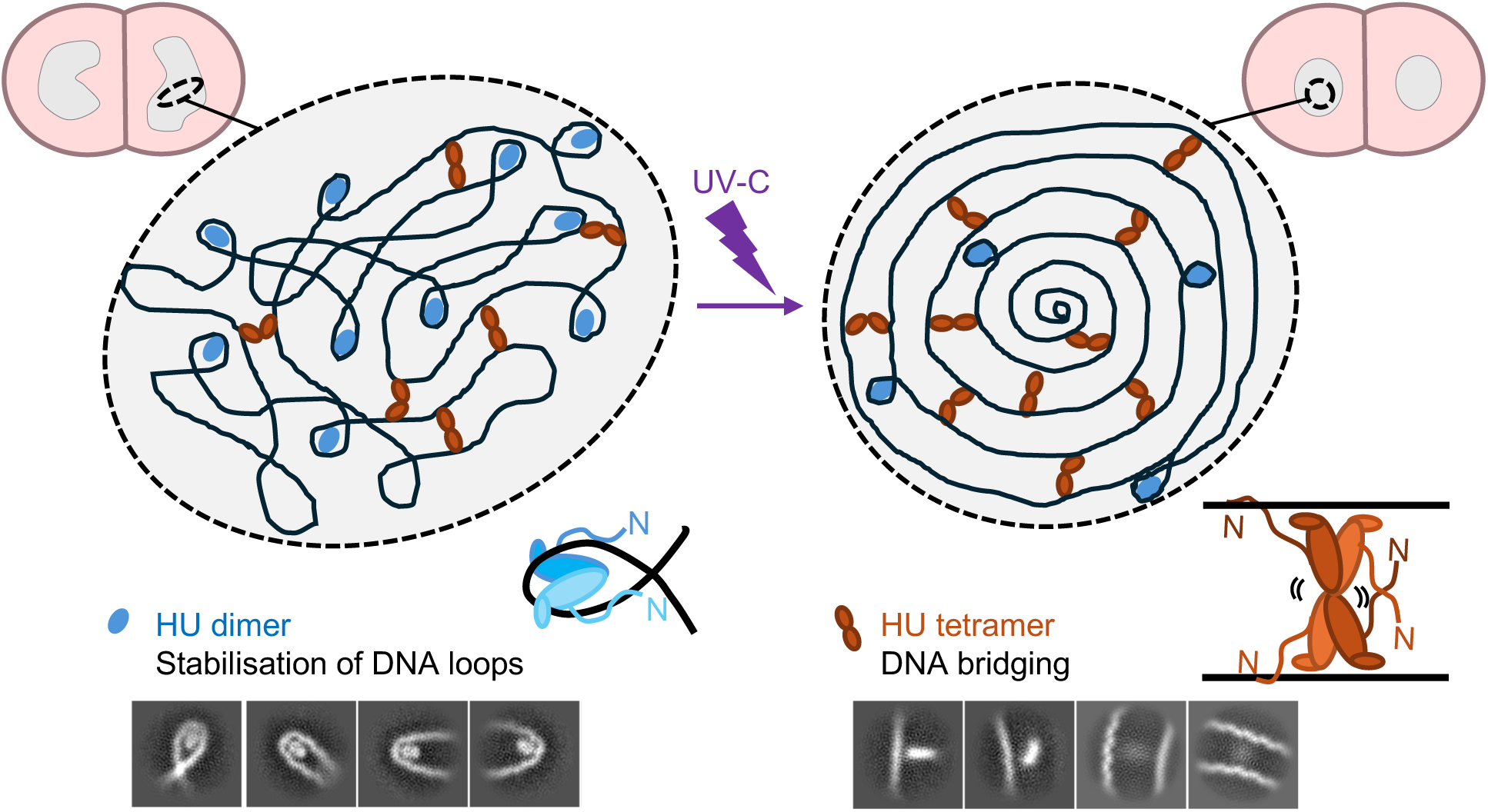
Schematic model of *Dr*HU’s versatile role in genome organization. Under normal growth conditions (left), the *D. radiodurans* nucleoid adopts a loosely organised DNA mesh with no obvious regular pattern. Exposure to UV-C (right) triggers nucleoid remodelling, reducing its size, increasing its sphericity, and inducing the formation of a blue liquid crystalline phase with a regular inter-DNA distance of ∼8.5 nm. Central to this process is *Dr*HU’s versatile DNA binding, which occurs through at least two distinct modes: dimers (blue) stabilise tight DNA bends via β-hairpin insertion into the minor groove, while tetramers (orange/brown)—comprising flexibly associated back-to-back dimers—bridge distant DNA filaments. This bridging is largely dependent on *Dr*HU’s disordered N-tails, which mediate non-specific DNA binding and promote tetramer formation. In cells, the 3D genome architecture and dynamics (illustrated in 2D here) likely result from a balance between these two complementary properties. Under normal growth conditions, most of *Dr*HU exists as dimers bound to tight DNA loops generated during transcription and replication, whereas under stress, an increased fraction of DNA-bridging *Dr*HU drives the transition from a loose DNA mesh to a blue liquid crystalline phase, which is more favourable for efficient genome repair.

One of the most striking observations from our study is the ability of *Dr*HU to drive supercoiled plasmid DNA into eight-shaped double spirals that tile the cryo-EM grid into a 2D lattice with a regular inter-filament spacing. To our knowledge, such a uniform, protein-mediated DNA lattice has not been previously described and provides the first direct visualisation of the ability of HU to drive higher-order organisation of a circular, supercoiled DNA molecule, through its DNA bridging capacity. We cannot exclude that blotting and air-water interface-related phenomena may also contribute to some extent to the formation of this remarkable 2D lattice, but several observations argue that this regular arrangement is largely driven by *D*rHU: (i) the arrangement occurs reproducibly in the presence of *Dr*HU only, and at a specific *Dr*HU:plasmid ratio, (ii) none of the *Dr*HU variants reproduce this arrangement, and (iii) the 2D patterns obtained are dependent on DNA topology: the double spiral assembly occurs only with supercoiled, while parallel DNA filaments were observed with both linear and supercoiled plasmids, suggesting that *Dr*HU exploits the mechanical stress of supercoiling to impose higher-order geometric constraints. The uniform spacing between neighbouring DNA filaments (∼9 nm) is a key feature of these assemblies, and was observed with both supercoiled and linear DNA, indicating that while supercoiling is required for spiral formation, the ability of *Dr*HU to set a uniform inter-filament distance is its intrinsic property, independent of DNA topology. Our mutational analysis demonstrates that the N-tail of *Dr*HU and its multivalent binding to DNA, but not the tight DNA binding mode mediated by the β-hairpin, is required for this key function in spatial organisation and ordering of the DNA. The two modes of DNA binding of *Dr*HU thus appear to play distinct physiological roles, and together enable *Dr*HU to stabilise tight DNA loops, bridge adjacent DNA molecules, and impose uniform inter-filament spacing, thereby linking molecular interactions to supramolecular genome organisation (Fig. 4). Furthermore, *Dr*HU’s capacity to assemble as both dimers and tetramers is certainly also an important determinant of its DNA bridging capacity by setting the spacing between parallel DNA filaments.

These findings resonate with recent *in vivo* studies revealing heterogeneous HU populations with distinct diffusion properties. HU has indeed been shown to exist in multiple dynamically exchanging states in bacterial cells, with some fractions tightly bound to DNA and others more freely diffusing, both of which contribute to the viscoelastic properties and fluidity of the bacterial nucleoid^45,16,13,46,17^. It is tempting to speculate that these two populations may correspond to HU engaged in high-affinity site-specific binding to distorted or tightly constrained DNA structures (β-hairpin-dependent) and to low-affinity, multivalent interactions of HU across the entire genome (N-tail-dependent), with the latter potentially facilitating rapid redistribution of HU along the chromosome to ensure cell-cycle dependent or stress-induced nucleoid remodelling and transcriptional reprogramming^47,16,48,43,17^.

To further explore this hypothesis and assess the functional relevance of our *in vitro* observations, we initiated an *in situ* cryo-ET analysis of *D. radiodurans* nucleoids. While under normal growth conditions, the nucleoid adopts a loosely organised DNA mesh exhibiting no obvious regular pattern, UV stress triggers its dramatic remodelling. Macroscopically, this is evident in the cryo-ET data as a marked change in nucleoid size and shape, consistent with our recent fluorescence microscopy study of UV-induced nucleoid remodelling^17^ (Fig. 4). At the molecular level, our tomograms reveal that the DNA in UV-irradiated bacteria forms a blue liquid crystalline phase, assembled from double-twist cylinders with an inter-DNA distance (∼8.5 nm) remarkably similar to that measured on the *in vitro Dr*HU-DNA double spirals. The close agreement between the measurements made on the 2D supramolecular assemblies observed *in vitro* and the 3D nucleoids of irradiated cells *in situ* strongly argues for shared underlying principles and the involvement of *Dr*HU, and its N-tail- and tetramerisation-dependent DNA bridging capacity, as a central player in DNA organisation across both contexts. This interpretation is further supported by our earlier single-particle tracking data that demonstrated a large increase in the fraction of mobile *Dr*HU (weakly bound to DNA) in cells exposed to UV-C light, which may favour the transition from a loose mesh to the liquid crystalline phase^17^.

Converging indirect evidence suggests that this blue liquid crystalline phase^7,32,33,49^ may serve multiple functional roles, and notably provide a physical framework that intrinsically favours efficient genome maintenance. The local alignment of DNA effectively reduces the dimensionality of lesion search from a random 3D search to quasi-1D scanning, while lowering the risk of aberrant recombination during repair, both of which would contribute to enhanced DNA repair efficiency and genome stability^50^. Orientational order further presents DNA in a uniform geometry, with grooves facing consistent directions over mesoscopic distances, minimising steric clashes and facilitating predictable docking and processive scanning. The inherent viscoelastic properties of dense yet fluid blue liquid crystalline phases and the characteristic geometric defects of such phases that locally relax packing and enhance dynamics^51^, may further enhance the accessibility of repair enzymes to damaged sites while maintaining global compaction^52^. This compromise between higher-order structure and dynamics may be critical for the extraordinary resistance of *D. radiodurans* to DNA damage.

In conclusion, this study establishes *Dr*HU as a versatile architect, capable of binding and protecting DNA loops, but also bridging and spacing DNA through distinct structural elements and oligomeric states. *Dr*HU’s multifunctionality likely compensates for the absence of additional NAPs in *D. radiodurans*, offering a compact, stress-responsive genome organisation system. By elucidating the structural and mechanistic principles that govern *Dr*HU-mediated higher-order chromatin organisation both *in vitro* and *in vivo*, this work significantly advances our understanding of bacterial chromatin architecture and NAP-driven genome remodeling in response to stress, and provides a solid framework for future endeavours.

## Methods

### Protein expression and purification

Cloning, expression, and purification of *D. radiodurans* HU has been described previously^42^. The N-terminally deleted (missing residues 1-28) form of HU (HU^Δ28^) was prepared by subcloning a restriction enzyme digested PCR fragment amplified with the following primers 5’ – AA CCA TGG GA GCC GAC AGC GGC AAG GTC GCC – 3’ and 5’ – TT AAG CTT TTA CAG GTT GCC CTT GAG GGT GC – 3’ (MWG Eurofins) into pProexHtB. The β-hairpin deletion mutants (HU^ΔHp^ and HU^Δ28ΔHp^) in either the wild-type or Δ28 constructs of HU were prepared by site-directed mutagenesis using the following primer pair: 5’-CTC TCG GTC AAG GAA ACC GGC AGC GGC AAG AAG GTC GCA TTC-3’ and 5’-GAA TGC GAC CTT CTT GCC GCT GCC GGT TTC CTT GAC CGA GAG-3’ (MWG Eurofins). The β-hairpin residues 89 to 106 were deleted and replaced by a glycine and serine residue. All clones were verified by DNA sequencing (Genewiz). Mutant HU constructs were expressed with cleavable (TEV protease) N-terminal His-tags in BL21 (DE3) at 20°C overnight and were purified as for the wild-type using a three step protocol involving an initial Nickel-affinity chromatography, followed by a heparin chromatography before cleavage of the His-tag, and a final size-exclusion chromatography (SEC) in 20 mM Tris-HCl, pH8.0, 100 mM NaCl^42^. Protein purity was verified by SDS-PAGE analysis. In the absence of aromatic residues, the concentration of HU proteins was determined by SDS-PAGE analysis using known concentrations of lysozyme (similar molecular weight) as a standard curve. Aliquots of concentrated HU (between 0.4 and 2 mM) were stored at -80°C and were thawed on ice before use.

### pUC19 DNA preparation

Plasmid pUC19 DNA was amplified in DH5α cells grown in LB supplemented with 100 μg.mL^−1^ ampicillin. Plasmid DNA was extracted from 200 mL overnight cultures using the NucleoBond Xtra Midi kit (Macherey-Nagel) following manufacturer’s instructions. The purified supercoiled plasmid DNA was resuspended in 200 μL sterile Milli-Q water, typically at a concentration between 2 and 4 μg.μL^−1^. The stock of supercoiled DNA was aliquoted and stored at −20°C. The quality of the preparation was verified by electrophoresis on a 0.7% TBE agarose gel. Linearized pUC19 plasmid DNA was prepared by digesting 36 μg supercoiled plasmid with EcoRI-HF (New England Biolabs) for 90 min at 37°C in 1xCutsmart buffer. Linear plasmid DNA were then loaded on a 0.7% TBE agarose gel, gel extracted using the Gel and PCR Clean Up kit (Macherey-Nagel) following manufacturer’s instructions and eluted in 2x30 µL at a concentration of ∼0.8 µg.µL^-1^. The conformation and purity of the linearised DNA were assessed by electrophoresis on a 1% TBE agarose gel followed by staining with Gel Red (Interchim) and visualization using a ChemiDoc MP imager (BioRad). Linear pUC19 DNA was aliquoted and stored at −20°C.

### Cryo-EM sample preparation

Purified pUC19 was diluted to 1 µM in H_2_O, and purified *Dr*HU^WT^ was diluted to 100 µM in SEC-Buffer (20 mM Tris-HCl, pH8.0, 100 mM NaCl) supplemented with 1 mM MgCl_2_. Prior to deposition on the cryo-EM grids, 5 µL of 1 µM pUC19 and 5 µl of 100 µM *Dr*HU^WT^ were mixed together with 15 µL of H_2_O. For cryo-EM grid preparation, a 3 µl droplet of the sample was applied to a glow-discharged Quantifoil R2/1 Cu 400 mesh grid and plunge frozen in liquid ethane using a Vitrobot Mark IV (FEI) operated at room temperature and 100% humidity. Grids were clipped and stored in liquid nitrogen for further use.

### Cryo-EM data acquisition

All cryo-EM images except the phase plate data set were acquired at the IBS Glacios microscope (Thermo Fisher Scientific) operated at 200 kV. Difficulties to locate *Dr*HU in these images likely arise from the small size of the *Dr*HU dimer, the dominance of the DNA signal and the fact that 2D classification is driven primarily by variations in DNA curvature and inter-filament spacing rather than by the presence, position or orientation of *Dr*HU. To increase contrast and ease of particle picking, a data set of 1503 Movies for SC-pUC19 mixed with *Dr*HU at a 1:100 ratio was collected at the Umeå Center for Electron Microscopy, on a Titan Krios (Thermo Fisher Scientific), equipped with a Volta phase plate and a Gatan K2 camera, and operated at 300 kV. Data was collected with a nominal magnification of 165000 x, resulting in a pixel size of 0.82 Å/px, a total dose of 53 e-/Å^2^ over 40 fractions and a defocus range of -0.5 µm to -1.2 µm.

### Cryo-EM data processing

Movies were imported in to CryoSPARC^53^, motion corrected with Patch Motion Correction, followed by CTF estimation using Patch CTF Estimation. After manually picking 322 particles, a first Topaz model for particle picking was trained^54^. Extracted particles were subjected to 2D classification, whereby one 2D class average showed a DNA loop that seemed to contain *Dr*HU. Particles of these classes were used to retrain Topaz and perform several consecutive repicking rounds. 2D class averaging resulted in 20,152 well-defined loop-containing classes featuring an extra density inside the DNA loop. These classes were subjected to heterogenous 3D refinement into 2 classes. The better-defined 3D class, with 11,324 particles, was subjected to a final 3D refinement resulting in a ∼14 Å resolution cryo-EM map. As with this approach most *Dr*HU particles outside the tight DNA loops remained unpicked, a fully manual picking was carried out in parallel. Manual picking revealed 2D class averages of other binding states along the DNA as shown in Fig. 1. However, higher alignment inaccuracies for these particles and the absence of sufficiently different orientations of the same type of particles made it impossible to achieve a meaningful 3D cryo-EM reconstruction.

### S1 and Bal31 Nuclease DNA digestion assays

S1 nuclease time course experiments were performed by incubating 20 nM supercoiled pUC19 alone or in the presence of 2 µM *Dr*HU (wild-type or mutants) with 1µL S1 nuclease (Takara) in 0.5X S1 buffer in a reaction volume of 75 µL. Reactions were run at 37°C and stopped at 0, 2, 15 and 30 min time points, by mixing 15 µL reaction with 2X DNA purple loading dye (New England Biolabs) and heating samples at 95°C for 90 sec. S1 and Bal31-treated samples were then analysed on 1% TBE agarose gel (electrophoresis at 100 V at 4°C for 50 min) followed by staining with Gel Red (Interchim) and visualisation using a ChemiDoc MP imager (BioRad). Bal31 digestion experiments were performed by incubating 100 nM supercoiled pUC19 alone or in the presence of 10 µM *Dr*HU (wild-type or mutants) with 1µL Bal31 nuclease (Takara) in salt adjusted buffer (20 mM Tris, 100 mM NaCl, 5 mM CaCl_2_, 5 mM MgCl_2_) in a reaction volume of 10 µL at 30°C for 0, 1 and 2 min. Reactions were stopped by adding binding buffer of PCR Clean Up kit. The digested DNA was then purified using the Gel and PCR Clean Up kit (Macherey-Nagel) following manufacturer’s instructions and eluted in 20µL elution buffer.

### DNA binding assays

Binding of HU (wild-type and mutants) to pUC19 plasmid DNA (linear or supercoiled) was assessed by electrophoretic mobility shift assay (EMSA). For this, 3 nM plasmid DNA was incubated with 0 to 10 µM HU in DNA binding buffer (20 mM Tris pH 8, 50 mM NaCl, 1 mM MgCl_2_, 0.02 mg/mL BSA) for 15 min at 30°C before loading on a 0.7% TBE agarose gel, which was run at 4°C for 1h at 100V. The DNA was then stained with Gel Red (Interchim) and visualized using a ChemiDoc MP imager (BioRad). Binding of HU (wild-type and mutants) to short dsDNA oligonucleotides (50 mer dsDNA) was assessed by fluorescence polarisation (FP) experiments. For this, a 50 mer dsDNA substrate was prepared by annealing a 6-FAM-labelled strand (5’- [FAM] GAC TAC GTA CTG TTA CGG CTC CAT C G CTA CCG CAA TCA GGC

CAG ATC TGC -3’; MWG Eurofins) with its complementary strand (5’- GCA GAT CTG GCC TGA TTG CGG TA GCG ATG GAG CCG TAA CAG TAC GTA GTC -3’; MWG Eurofins) in 20 mM Tris pH8, 50 mM NaCl and 0.5 mM EDTA. FP reactions were prepared by mixing 2 nM 50 mer FAM-dsDNA with 0 to 40 µM HU in DNA binding buffer in a final reaction volume of 40 µL. Equilibrium FP measurements were performed on a Clariostar microplate reader (BMG Labtech), fitted with polarization filters using an excitation wavelength of 482-16 nm and an emission of 530-40 nm. In all cases, after subtracting the polarization values obtained for DNA alone, the mean data from at least three independent experiments were fitted to either a one-site specific binding model with Hill slope (Y = Bmax*X^h^/(Kd^h^+X^h^) or a two-site specific model (Y = [BmaxHi*X/(KdHi+X)] + [BmaxLo*X/(KdLo+X)]) using GraphPad Prism 10, where Bmax is the difference between the polarization of completely bound and completely free oligo, h is the Hill slope, X is the HU concentration and Kd is the equilibrium dissociation constant. All fits were very good with R2 values above 0.96.

### BS3 Chemical crosslinking

After a buffer exchange step into 20 mM Na-phosphate pH 8, 100 mM NaCl, 50 µM wild-type and mutant *Dr*HU were chemically crosslinked with 0-5 mM bis(sulfosuccinimidyl)suberate (BS3) for 30 min at 25°C before separation on a 10% SDS-PAGE gel (BioRad) stained with Instant Blue to reveal the protein bands. Gel images were acquired and quantified using a ChemiDoc MP imager and ImageLab software (BioRad).

### Cryo-ET sample preparation

Exponentially growing (OD650 ∼0.5) *D. radiodurans* R1 cultures (50 mL/sample) were prepared by serially diluting a preculture in 50 mL Tryptone-Glucose-Yeast extract (TGY) 2X medium and growing the cells at 30°C overnight. Non-irradiated cells were pelleted by centrifugation at 5000 rcf for 10 min and resuspended in 0.5 mL TGY2X, flash-frozen in liquid nitrogen and stored at - 80°C. To prepare irradiated samples, cells were pelleted and resuspended in 10 mL minimal (M9-DR) medium and irradiated with UV-C light in a 10 x10 cm petri dish at a dose of 1.9kJ/m2 using a Stratalinker UV oven, as described previously^17^. Immediately after irradiation, cells were returned to TGY2X medium and placed at 30°C in a shaking incubator to recover for 1h. After this, cultures were pelleted as described above, resuspended in 0.5 mL TGY2X, flash-frozen in liquid nitrogen and stored at -80°C until further use. To prepare EM grids, 0.5 mL aliquots corresponding to 50 mL cultures of either non-irradiated or irradiated samples were thawed on ice and 250 µL of the cell suspension was harvested at 2000 rcf for 3 min, after which cells were washed 3× with 500 µL phosphate buffered saline (PBS). The resulting cell pellet was initially 4X diluted, from which a dilution series was prepared. Different dilutions were deposited on an EM-grid, dry blotted and observed under a light microscope to check for optimal concentration and sample spread. Quantifoil R2/2, 200 mesh, Cu grids were then used for plunge freezing in liquid ethane using a Leica GP2 under the following conditions (Temperature: 20°C, 99 % humidity, Blot time 3-4 s, Volume: 4 µl). Vitrified grids were screened on a Zeiss LSM900 Airyscan2 cryo-fluorescence light microsocpe equipped with a Linkam cryostage to find optimal concentration and cell distribution for focused ion beam (FIB) milling.

### Focused Ion Beam milling

Cryo-FIB milling was carried out on an Aquilos 2 cryo-FIB/SEM instrument (ThermoFisher) using AutoTEM (ThermoFisher). The samples were sputter coated with and inorganic platinum for 30s (30 mA, 0.1 mbar), after which an organometallic platinum layer was applied using the integrated gas injection system (GIS) for 1.5 - 2 min, followed by an additional round of platinum sputtering for 30 s using the same parameters. Stress relief cuts of 300 nm width, 10 µm height and 5 µm distance to the lamellae were introduced to improve lamellae stability. Lamellae were milled to a thickness of ∼180 nm with a width of 10 - 16 µm using a milling angle of 7°. Milling was carried out in three steps with decreasing ion beam currents of 1 nA, 0.3 nA and 0.1 nA followed by two polishing steps and 50 pA and 30 pA to reach ∼180 nm thickness. After AutoTEM milling finished, lamellae were post-polished manually by milling the top of the lamellae (above the GIS layer), with a rectangular pattern top-bottom scan direction at 30 pA tilting up by 0.5° to reach the back of the lamellae, which makes lamellae thickness more homogeneous. A final sputter coating with platinum was applied for 3 s (30 mA, 0.1 mbar).

### Cryo-ET data acquisition

Cryo-ET data collection was carried out on a Titan Krios Transmission Electron Microscope (ThermoFisher Scientific) equipped with a Falcon4 camera and operated 300 keV. The tilt series were acquired at 64 000× magnification with a pixel size of 1.968 Å/px, a defocus range between -2.0 and -5 µm and an approximate electron dose of per tilt image of 3.17 e-/Å^2^ for the non-irradiated *D. radiodurans* cells and 3.65 e-/Å^2^ for the irradiated *D. radiodurans* cells. Data was collected with SerialEM^55^ and PACE-tomo^56^ using a dose symmetric acquisition scheme with a tilt range between +54° and -66° in 3° increments.

### Cryo-ET data processing

Tilt images were motion corrected and CTF estimated by WarpTools version 2.0.0 (fs_ctf, c-grid 2x2x1)^57^. Bad tilt images were removed manually, followed by tilt series alignment using AreTomo2 version 1.1.3^58^. Alignments were imported back to WarpTools and CTF model was checked for correct handedness. Tilt series were then reconstructed with a 4x-binning factor to a resolution of 7.872 Å. Distance measurements between DNA filaments in IR *D. radiodurans* cells was done on single Z-slices using 3dmod^59^.

### Orientation field calculation and analysis

Orientation fields were computed from local image intensity variations to provide an additional means of evaluating and illustrating the organisation of the DNA filaments within the swirls (Supplementary Fig. S10). To minimise missing wedge artefacts, calculations were performed on averaged 5 nm thick central slices oriented perpendicular to the beam rather than on the 3D tomographic volumes. For each pixel, directional derivative responses were calculated using four 3×3 Sobel filters (0°, 45°, 90° and 135°). Filament orientation was defined as orthogonal to the dominant response direction. The calculated orientations were then locally averaged within circular patches of ∼40 nm diameter to reduce noise and to estimate the locally predominant orientation. A confidence score was computed for each pixel to differentiate regions containing well-aligned DNA filaments from those lacking a dominant orientation. This score is thus defined as the ratio of the response of the dominant directional derivative to the sum of responses across all considered directions, providing a measure of the reliability of the estimated orientation. As shown in Supplementary Fig. S10, the resulting orientation fields exhibit a clear spiral organisation, with high confidence (red) in regions containing DNA swirls and lower confidence (green/blue) towards the centres of the spirals and in regions with less ordered DNA.

### MD simulations

Initial structural models of wild-type (WT) and N-tail-truncated (Δ28) tetrameric *Dr*HU, in which each constituent dimer is complexed with a 36 mer dsDNA duplex, were constructed using the AlphaFold3 (AF3) web server^60^ (Supplementary Figure S7c). Although the *Dr*HU-DNA conformation predicted by AF3 in which DNA is wrapped around two *Dr*HU dimers is biased by available models in the database and is unlikely to be the conformation underlying the 2D spiral arrangement and 3D genome organisation, it provided a robust starting model for exploring dimer-dimer contacts underlying tetramer formation and the role of N-tails in both DNA binding and tetramer stabilisation by MD simulations. Four distinct structural ensembles were defined for the simulations: (i) *Dr*HU^WT^ bound to dsDNA, (ii) *Dr*HU^Δ28^ bound to dsDNA, (iii) apo *Dr*HU^WT^ and (iv) apo *Dr*HU^Δ28^ alone. The apo states were prepared simply by removing the bound DNA segments. All MD simulations were prepared using the AMBER tleap prgramme^61^. Each system was solvated in an octahedral box of water molecules (maintaining a minimum distance of 12 Å between the solute atoms and the box edge). Chemical neutralization and an ionic strength of 0.15 M were achieved by adding NaCl counterions. The ff14SB force field was used for protein atoms^62^, BSC1 for DNA segments^63^, while the water molecules were treated using the standard TIP3P model^64^. The simulation runs were carried out using NAMD3^65^ under full GPU acceleration. By leveraging the HMR protocol^66^, the equations of motion were integrated with an accelerated 4 fs time step under periodic boundary conditions. Non-bonded interactions were managed using a real-space cutoff of 8.0 Å coupled with a pair-list distance of 10.0 Å, and the Lennard-Jones parameters were updated using a long-range analytical correction. Long-range electrostatic interactions were computed via the Particle Mesh Ewald method^67^, using a grid spacing of 1.0 Å and an interpolation order of 4. Thermodynamic ensembles were maintained at a target temperature of 300 K using a Langevin thermostat. Concurrently, pressure was regulated at 1 atm using a Langevin piston Nosé-Hoover barostat^68^. To ensure geometric stability, all covalent bonds involving hydrogen atoms were strictly constrained using the SHAKE^69^ and RATTLE^70^ algorithms. To prevent structural distortion during initialization, a rigorous multi-step equilibration protocol was enforced to gradually relax the systems. First, an initial minimization and relaxation stage was conducted, consisting of 2,000 steps of energy minimization followed by a 10 ns equilibration window. During this phase, the heavy atoms of both the protein and the nucleic acids were restrained using a harmonic potential of 1 kcal mol^−1^ Å^−2^. Second, a backbone relaxation phase was initiated by releasing the restraints for all side-chain atoms; the system was then simulated for another 10 ns with harmonic restraints applied exclusively to the nucleic acid heavy atoms and the protein backbone heavy atoms. Third, a constraint scaling step was executed, wherein the remaining harmonic restraints were gradually downscaled from 1.0 to 0.1 kcal mol^−1^ Å^−2^ across five successive 2 ns windows. Following this final relaxation stage, an unrestrained production run of 1.03 μs was carried out for each of the four independent replicas per system, yielding a cumulative equilibrium sampling trajectory of 4 μs per ensemble for subsequent statistical analysis.

### MD trajectory analyses

To evaluate the structural stability and conformational drift of the complexes during the MD simulations, root-mean-square deviation (RMSD) trajectories were calculated using the MDTraj python library^71^. For each simulated system, the initial frame of the replicate trajectory served as the reference structure. Protein RMSD calculations were restricted to the backbone atoms (C, Cα, N, and O) excluding the flexible residues of the N-tails. For systems containing nucleic acids, DNA RMSD was computed across all heavy atoms. To extract representative conformations of the simulated complexes, geometric clustering of the MD trajectories was performed using RCDPeaks^72^. Clustering was executed for each ensemble of replicas per system with a local density distance cutoff (dc) set to 4 Å. The atom selection was restricted to the protein backbone atoms (C, Cα, N, and O) excluding the flexible residues of the N-tails. To monitor the distance separating the opposing DNA segments in tetrameric *Dr*HU during the simulations, the inter-DNA distance was monitored as a function of time. Specific DNA selection strings corresponding to the binding regions of the hairpins for Dimer-A and Dimer-B were dynamically assigned using a 3.0 Å contact proximity threshold. For each simulation frame, the target coordinates were updated, and the center of mass was calculated independently for each DNA selection segment. To monitor the persistence of specific interactions across the simulated ensembles, lysine contact trajectories were analyzed as a function of simulation time. Inter-dimer distances were computed frame-by-frame between the sidechain amino nitrogen (NZ) atoms of all selected lysine residues belonging to the designated dimeric segments (Dimer 1 and Dimer 2). Pairwise contacts were scored dynamically using a cutoff distance of 15 Å. Occupancy metrics were then derived as the average time a given pair remained below this spatial threshold throughout the entire production trajectory (all replicates merged).

## Data availability

Data supporting the findings of this study are available in the article and Supplementary Data. The phase plate data set and the low resolution cryo-EM map of *Dr*HU in a tight DNA loop will be deposited in EMPIAR and EMDB, respectively, upon publication.

## Acknowledgements

The Institut de Biologie Structurale (IBS) acknowledges integration into the Interdisciplinary Research Institute of Grenoble. The work benefitted from access to the Grenoble Instruct-ERIC center (ISBG; UAR 3518 CNRS-CEA-UGA-EMBL) within the Grenoble Partnership for Structural Biology, supported by the French Infrastructure for Integrated Structural Biology (ANR-10-INBS-0005-02) and the Grenoble Alliance for Integrated Structural & Cell Biology Labex, a project of the University Grenoble Alpes graduate school (Ecoles Universitaires de Recherche) CBH-EUR-GS (ANR-17-EURE-0003). For cryo-EM grid preparation and screening, this work used the EM platform of the Grenoble Instruct Center (ISBG; UMS 3518 CNRS-CEA-UJF-EMBL) with support from FRISBI (ANR-10-INSB-05-02) and GRAL (ANR-10-LABX-49-01) within the Grenoble Partnership for Structural Biology (PSB). The IBS EM facility is supported by the Rhône-Alpes Region, the Fondation Recherche Medicale (FRM), the fonds FEDER and the GIS-Infrastructures en Biologie Sante et Agronomie (IBISA). We are grateful to Dr. G. Schoehn for establishing and managing the IBS cryo-electron microscopy platform and for providing training and support, Dr. E. Zarkadas for assistance at the Glacios microscope, and A. Peuch for maintaining the joint IBS/EMBL computer cluster. The phase plate single particle cryo-EM data set was collected at the Umeå Center for Electron Microscopy (UCEM), the Umeå Cryo-EM node of SciLifeLab national research infrastructure. We thank the UCEM staff for training and support. We also acknowledge the access and services provided by the Instruct-ERIC Imaging Centre at the European Molecular Biology Laboratory (EMBL IC), generously supported by the Boehringer Ingelheim Foundation.

