## Supplemental Figures S1-S10 for "*Deinococcus radiodurans* HU: a versatile architect of nucleoid structure and plasticity"

**Supplementary Figures S1-S10**

**Supplementary movie legends**

**Supplementary movie 1:** Consecutive tomographic slices through a non-irradiated *D. radiodurans* cell, showing a nucleoid mesh in the bottom right. Slice thickness: 7.872 Å

**Supplementary movie 2:** Consecutive tomographic slices through an UV-C-irradiated *D. radiodurans* cell, showing DNA swirls in the nucleoid mesh. Slice thickness: 7.872 Å

**Supplementary movie 3:** Consecutive tomographic slices through an UV-C-irradiated *D. radiodurans* cell, showing a close-up view of a nucleoid in a double-twist cylinder arrangement, in which DNA filaments are aligned parallel to the cylinder axis at the centre, but progressively tilt outward with increasing radial distance, giving rise to a continuous, azimuthally symmetric twist in the cross-section. Slice thickness: 7.872 Å. Blue tracing along the DNA density facilitates the visualisation of the DNA swirls.

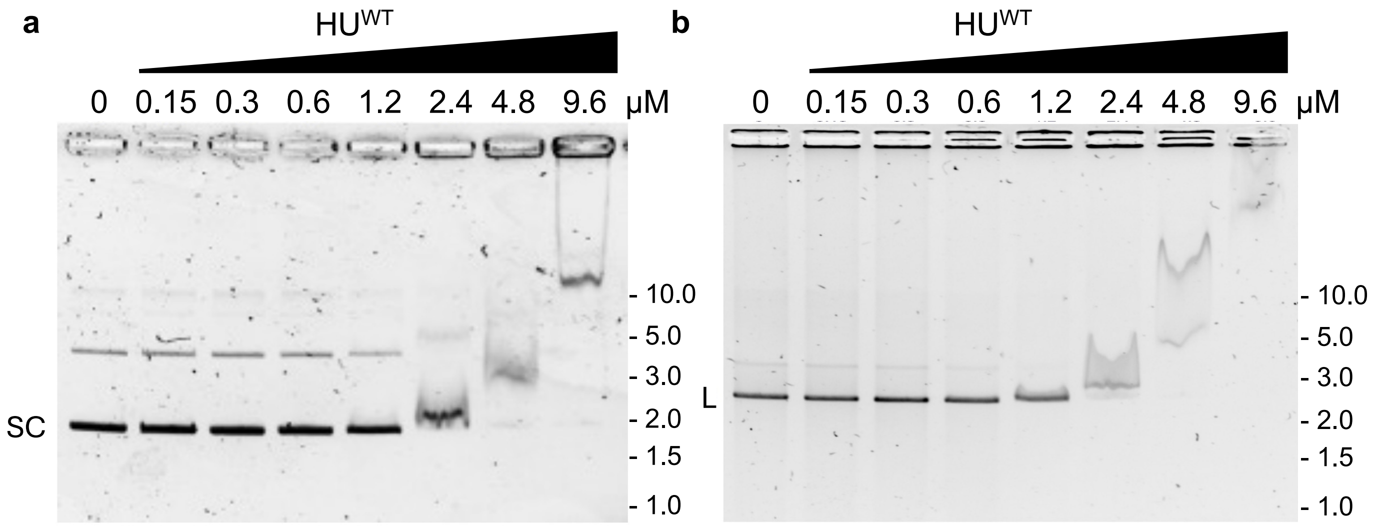


**Figure S1:** EMSA analysis of *Dr*HU binding to supercoiled and linear pUC19 plasmid. **(a)** $HU^{WT}$ (0 to 9.6 μM) binding to 3 nM supercoiled (SC) pUC19. **(b)** $HU^{WT}$ (0 to 9.6 μM) binding to 3 nM linear (L) pUC19. Molecular weight markers are indicated on the right of each gel in kb.

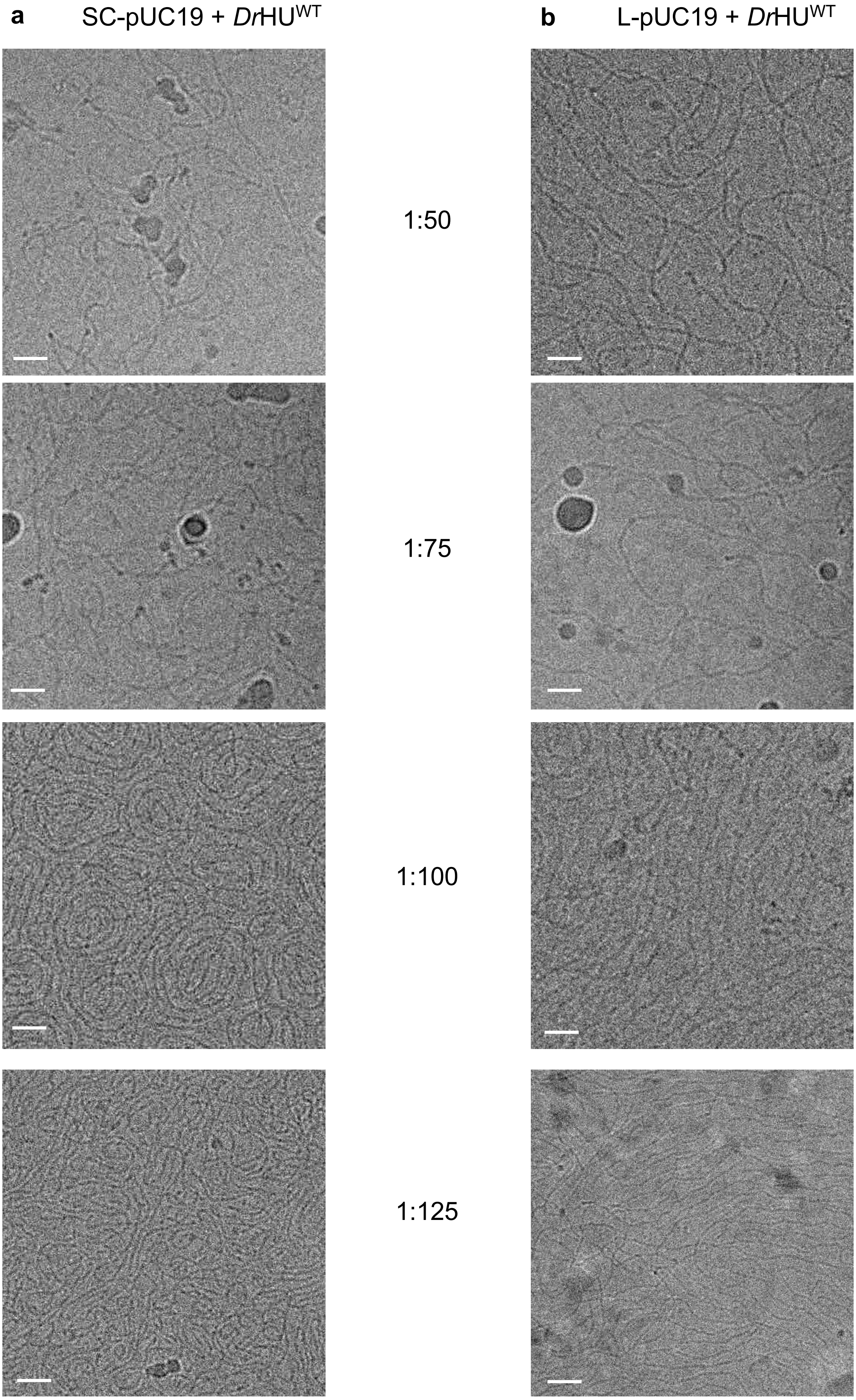


**Figure S2:** Representative cryo-EM micrographs of four different pUC19:*Dr*HU ratios: 1:50, 1:75, 1:100 and 1:125. **(a)** *Dr*HU$^{WT}$ assembled on supercoiled (SC) pUC19 DNA. **(b)** *Dr*HU$^{WT}$ assembled on linear (L) pUC19 DNA. Scale bar: 20 nm.

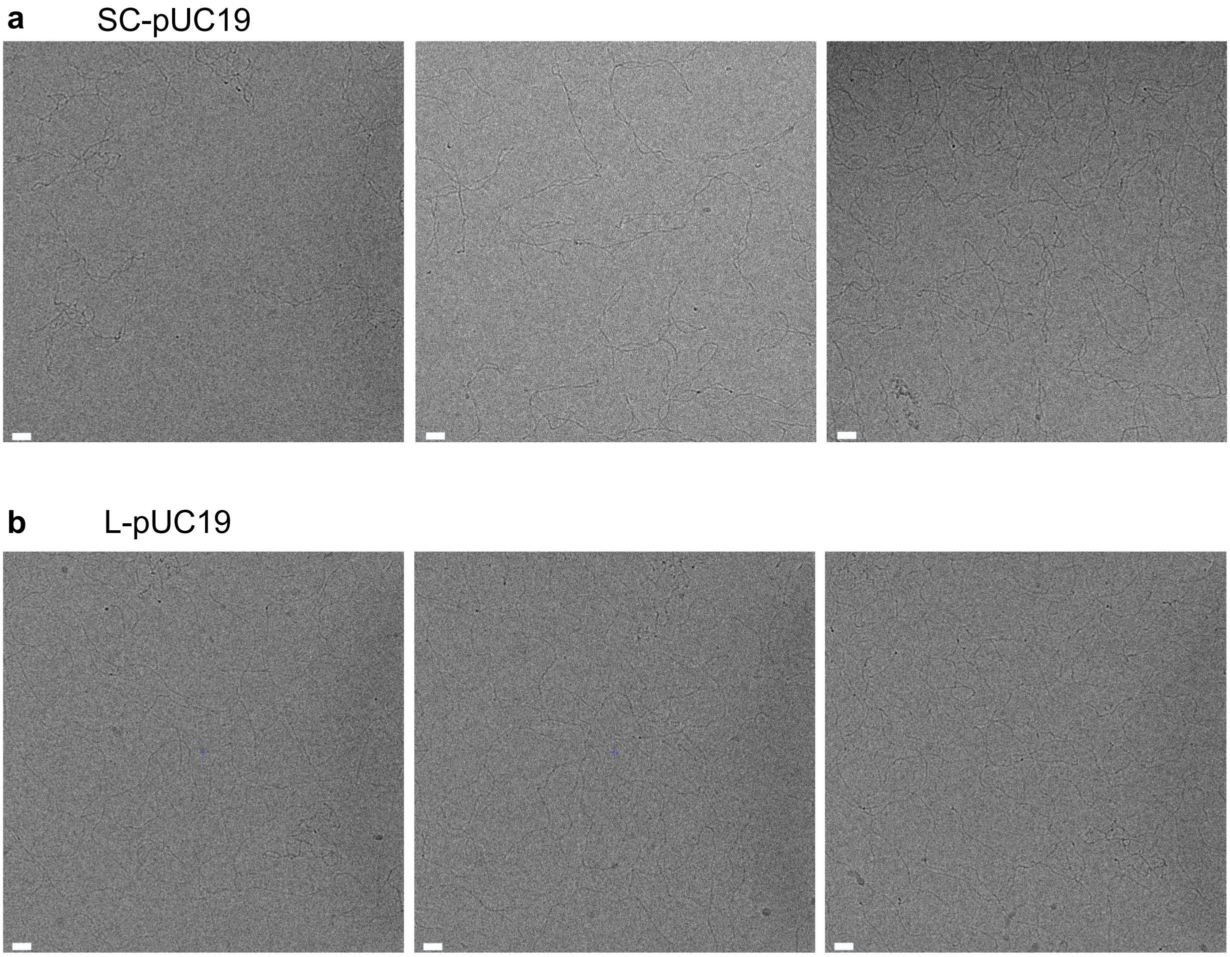


**Figure S3:** Representative cryo-EM micrographs of supercoiled (SC) pUC19 **(a)** and linear pUC19 **(b)**. Scale bar: 20 nm.

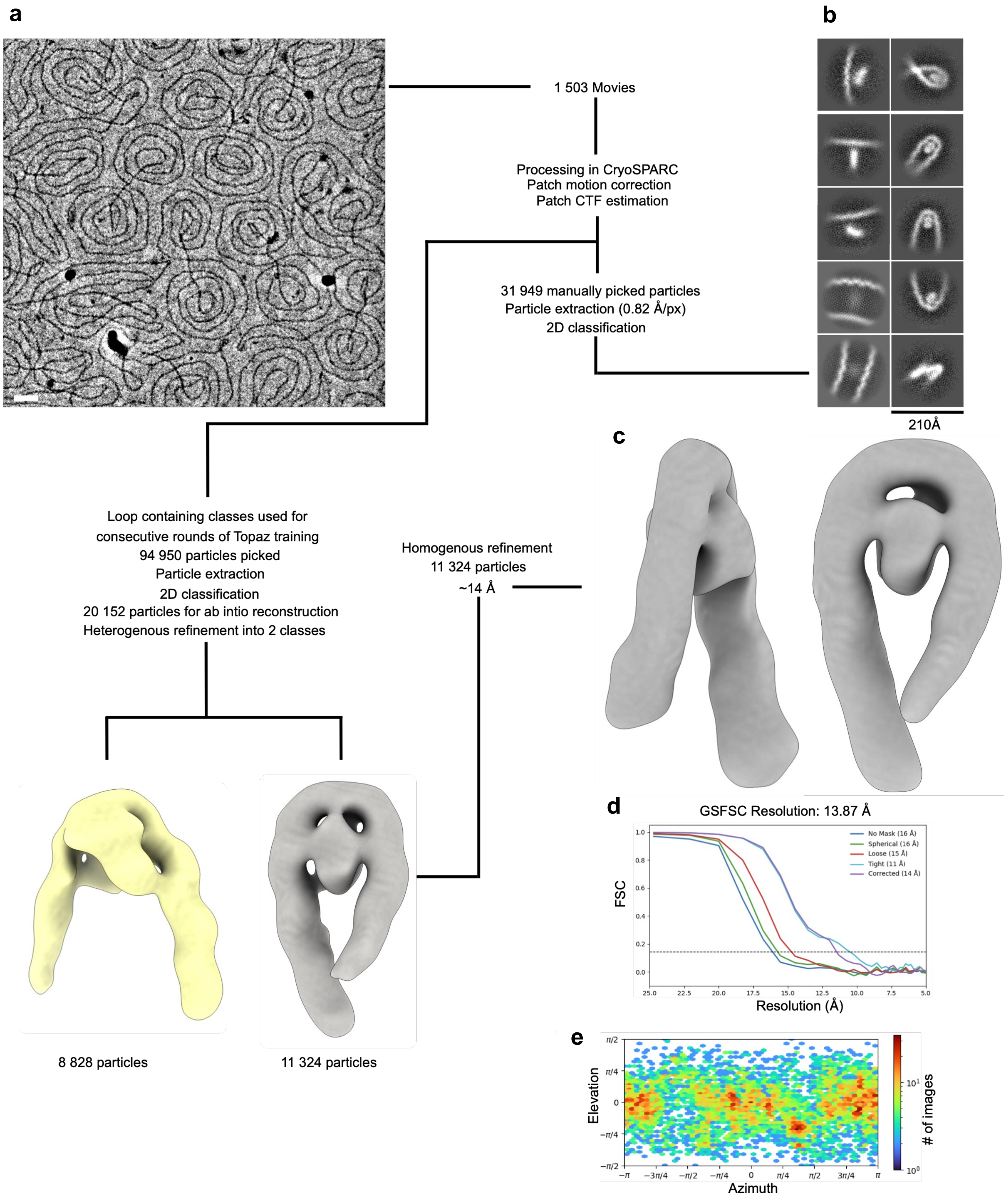


**Figure S4: (a)** Representative micrograph of SC-pUC19 + *Dr*HU$^{WT}$ (top left) and overview of the cryo-EM data processing pipeline. Scale bar: 20 nm. (**b)** Representative 2D class averages from manually picked particles. **(c)** Final map from CryoSPARC homogeneous 3D refinement. **(d)** CryoSPARC gold standard Fourier Shell Correlation with a resolution estimate of 14 Å at a cutoff of 0.143. **(e)** Particle orientations and angular distribution of the final reconstruction in CryoSPARC.

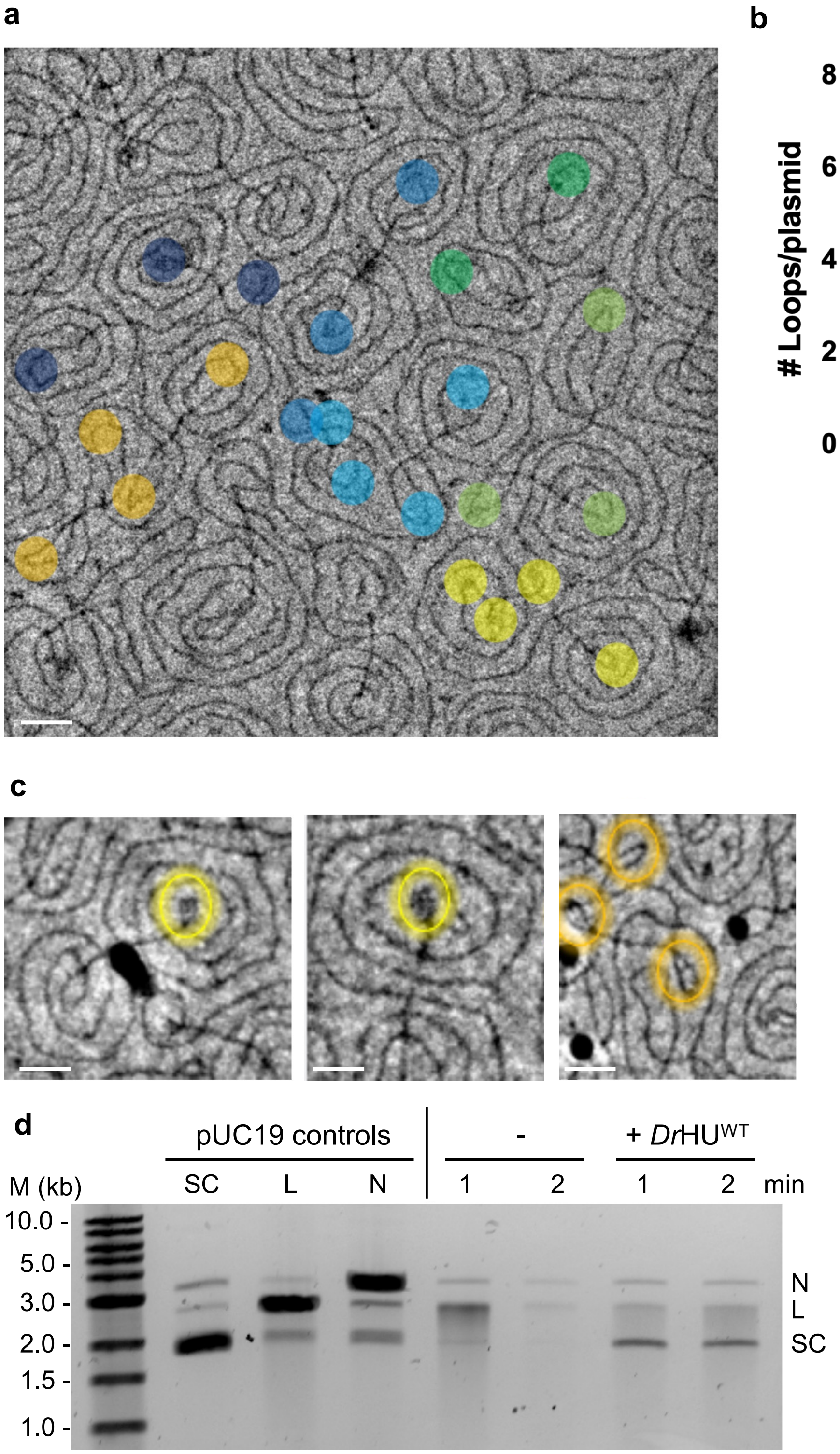


**Figure S5:** Analysis of tight loops in SC-pUC19-*Dr*HU micrographs. **(a)** Loop count in individual pUC19 molecules highlighted in different colors. Scale bar: 20 nm. **(b)** Box plot representation of the mean number of loops per plasmid (N=153). Error bars represent min and max values. **(c)** Close-up view of representative loop structures. Scale bars: 20 nm. **(d)** Agarose gel electrophoresis analysis of Bal31 nuclease assay, alongside supercoiled (SC), linear (L) and nicked (N) pUC19 plasmid, as indicated. SC-pUC19 alone was treated with Bal31 nuclease in the absence (lanes 5 and 6) or presence (lanes 7 and 8) of *Dr*HU$^{WT}$ for 1 and 2 min before analysis on gel. Nicked DNA was prepared by treating pUC19 with Nt.BspQI. Molecular weight markers are indicated in kb.

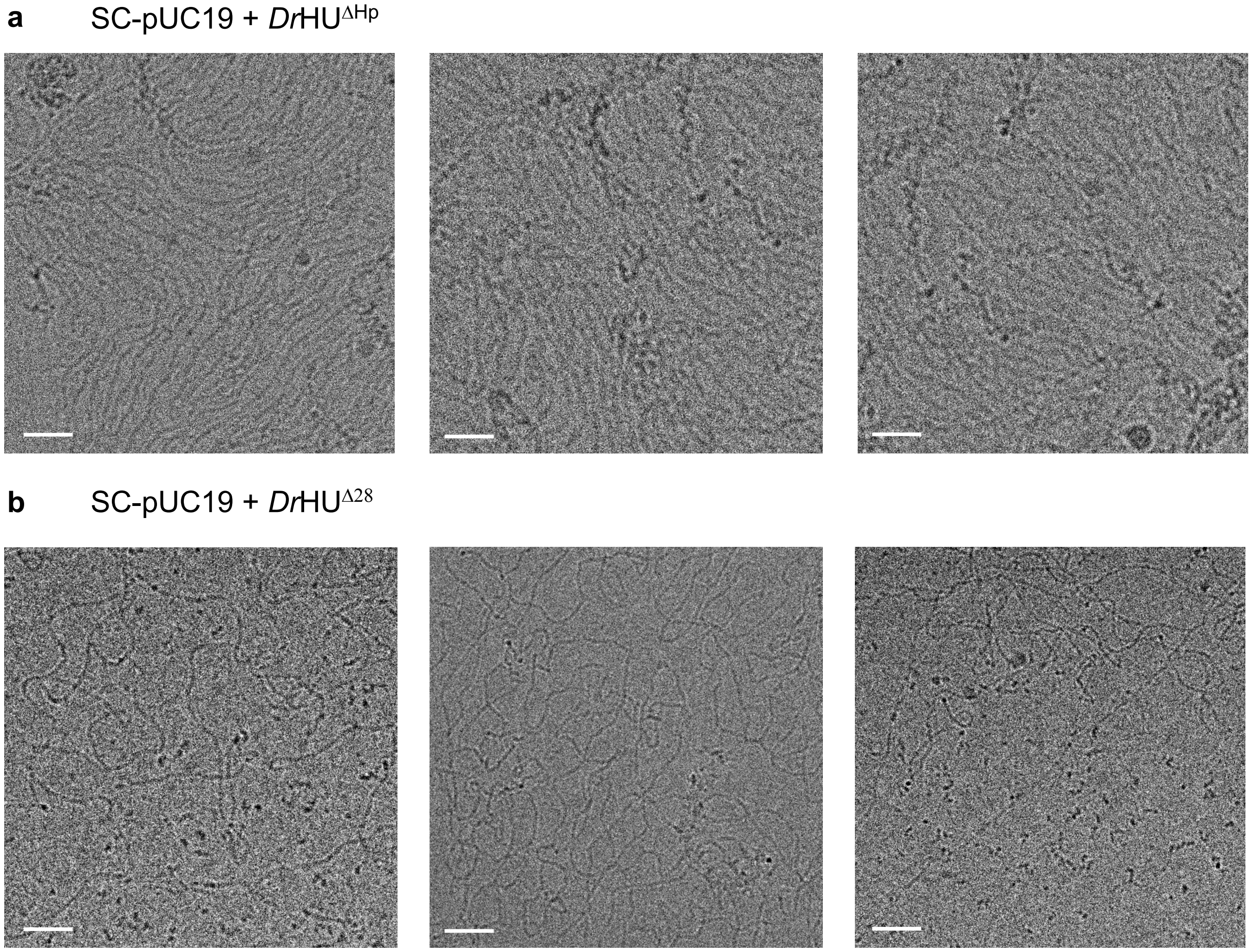


**Figure S6:** Representative cryo-EM micrographs of supercoiled (SC) pUC19 in the presence of either *Dr*HU$^{\Delta Hp}$ **(a)** or *Dr*HU$^{\Delta 28}$ **(b)**. Scale bar: 20 nm.

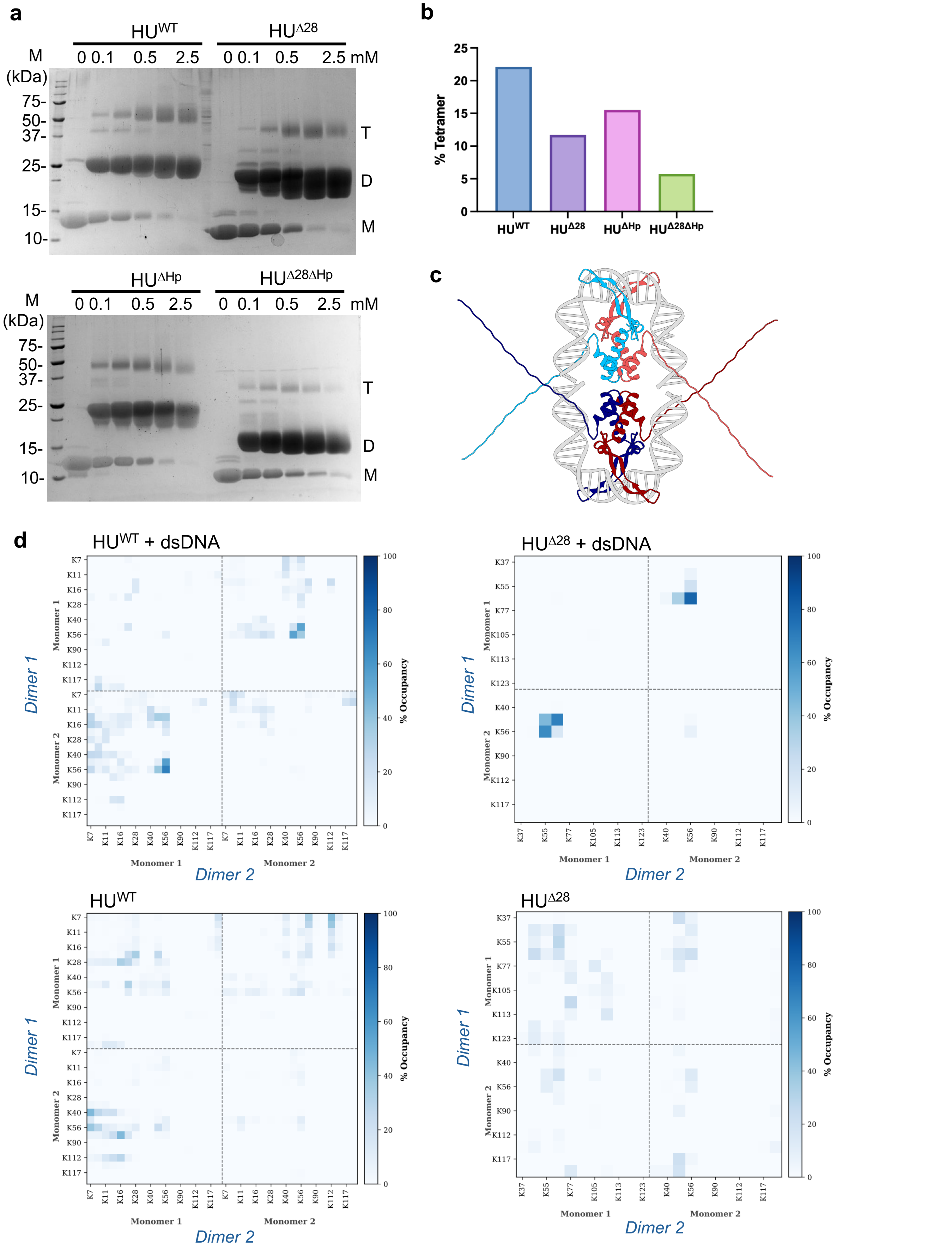


**Figure S7:** Chemical crosslinking of *Dr*HU and evidence for a tetrameric form. **(a)** SDS-PAGE analysis of BS3 chemical crosslinking (0-2.5 mM BS3) of wild-type (WT) and mutant *Dr*HU. Monomeric (M), dimeric (D) and tetrameric (T) forms of *Dr*HU can be seen. Molecular weight markers are indicated in kDa. **(b)** Gel-based quantification of the fraction of of WT and mutant *Dr*HU tetramer formed in the presence of 1 mM BS3 crosslinker. **(c)** AlphaFold3 generated model of tetrameric *Dr*HU bound to two 36bp DNA duplexes (see Methods for details) used as a starting model for MD simulations. One dimer is coloured in light blue and red, and the other in dark blue and red **(d)** Comprehensive raw matrices tracking individual pairwise Lys-Lys contact occupancy (%) between Dimer 1 (y-axis) and Dimer 2 (x-axis) using a cutoff distance of 15 Å during the various MD simulations. The colour gradient indicates contact persistence ranging from 0% to 100% occupancy.

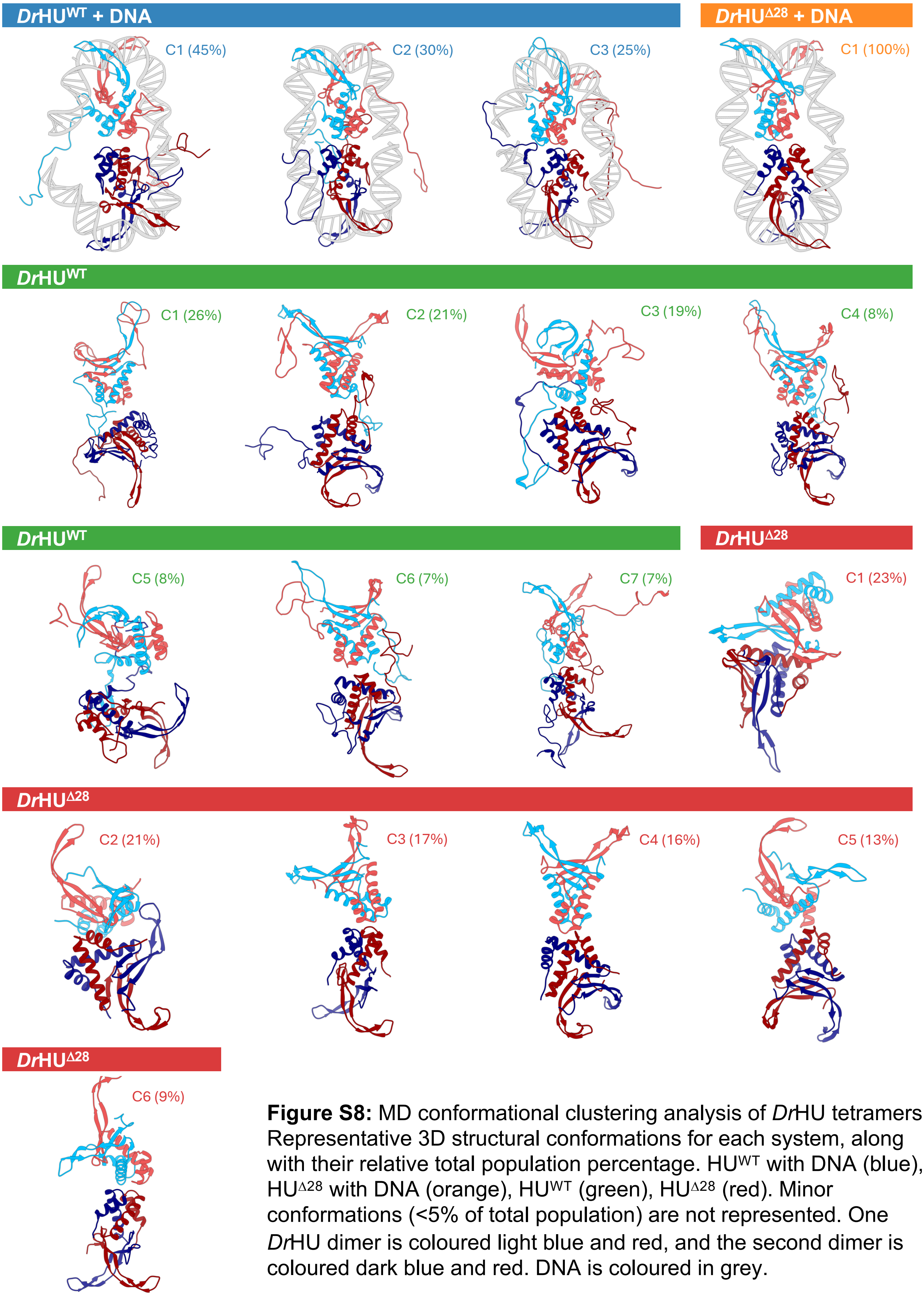


**Figure S8:** MD conformational clustering analysis of *Dr*HU tetramers. Representative 3D structural conformations for each system, along with their relative total population percentage. $HU^{WT}$ with DNA (blue), $HU^{\Delta 28}$ with DNA (orange), $HU^{WT}$ (green), $HU^{\Delta 28}$ (red). Minor conformations (<5% of total population) are not represented. One *Dr*HU dimer is coloured light blue and red, and the second dimer is coloured dark blue and red. DNA is coloured in grey.

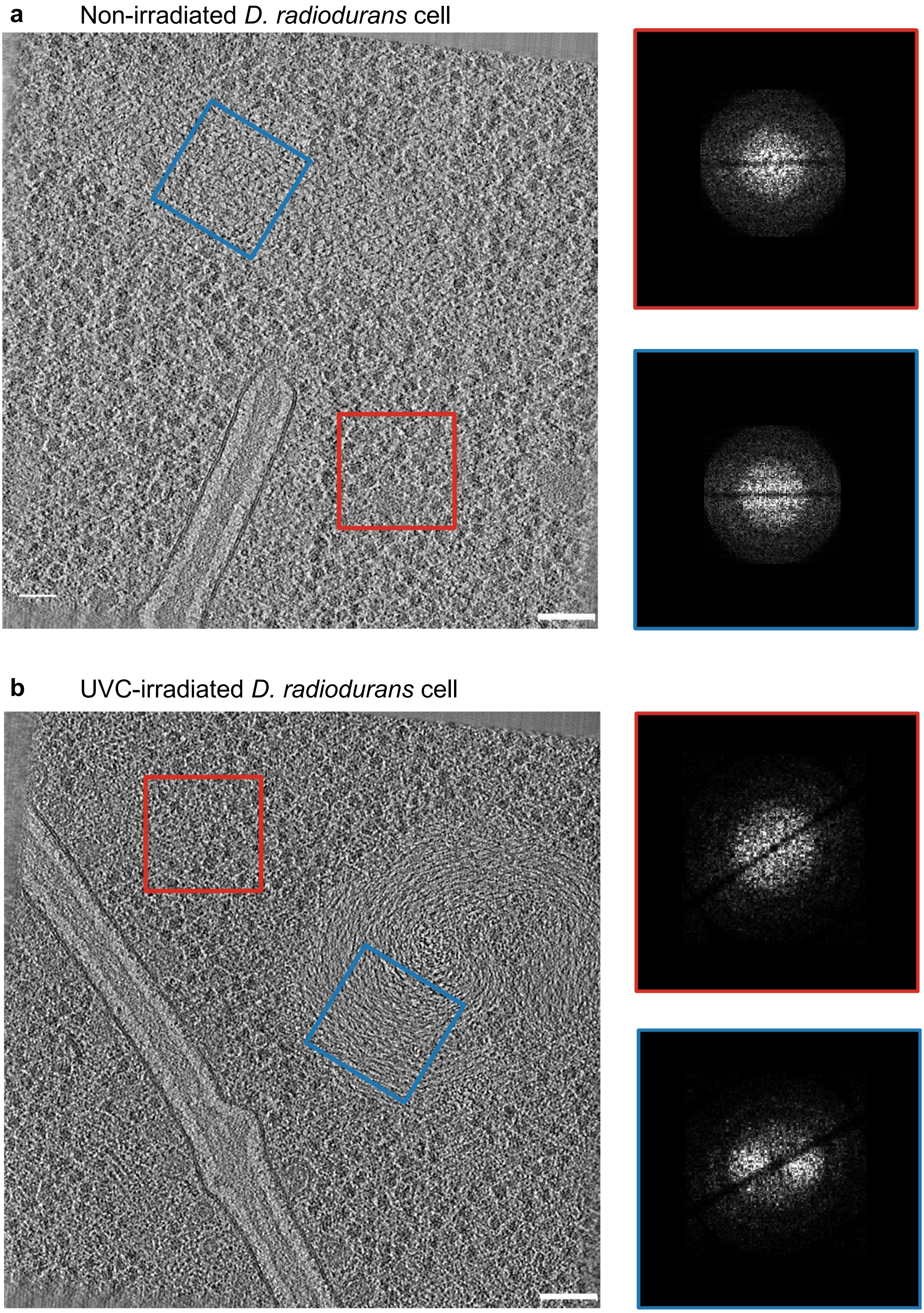


**Figure S9:** Slices through a representative tomogram of **(a)** non-irradiated and **(b)** UV-C irradiated *D. radiodurans* cells, illustrating distinct nucleoid morphologies and ultrastructure. Scale bar: 75 nm. Red and blue boxes on the left highlight a cytosolic and a nucleoid-containing region, respectively. The corresponding power spectra (right panels), calculated with the imod helixboxer tool and contoured in red or blue, indicate the presence of short-range positional order in the nucleoid of the irradiated cell (**b**, blue), in contrast to the non-irradiated nucleoid (**a**, blue) and the cytoplasmic regions (**a** and **b**, red). The broad peak in the Fourier transform of the irradiated nucleoid (**b**, blue) reflects the characteristic spacing of locally aligned DNA filaments. Together with the spiralling orientation fields (Fig. 3e,f and Extended Data Fig. 10), these observations suggest that after UV-C irradiation the nucleoid adopts a blue liquid-crystalline organisation.

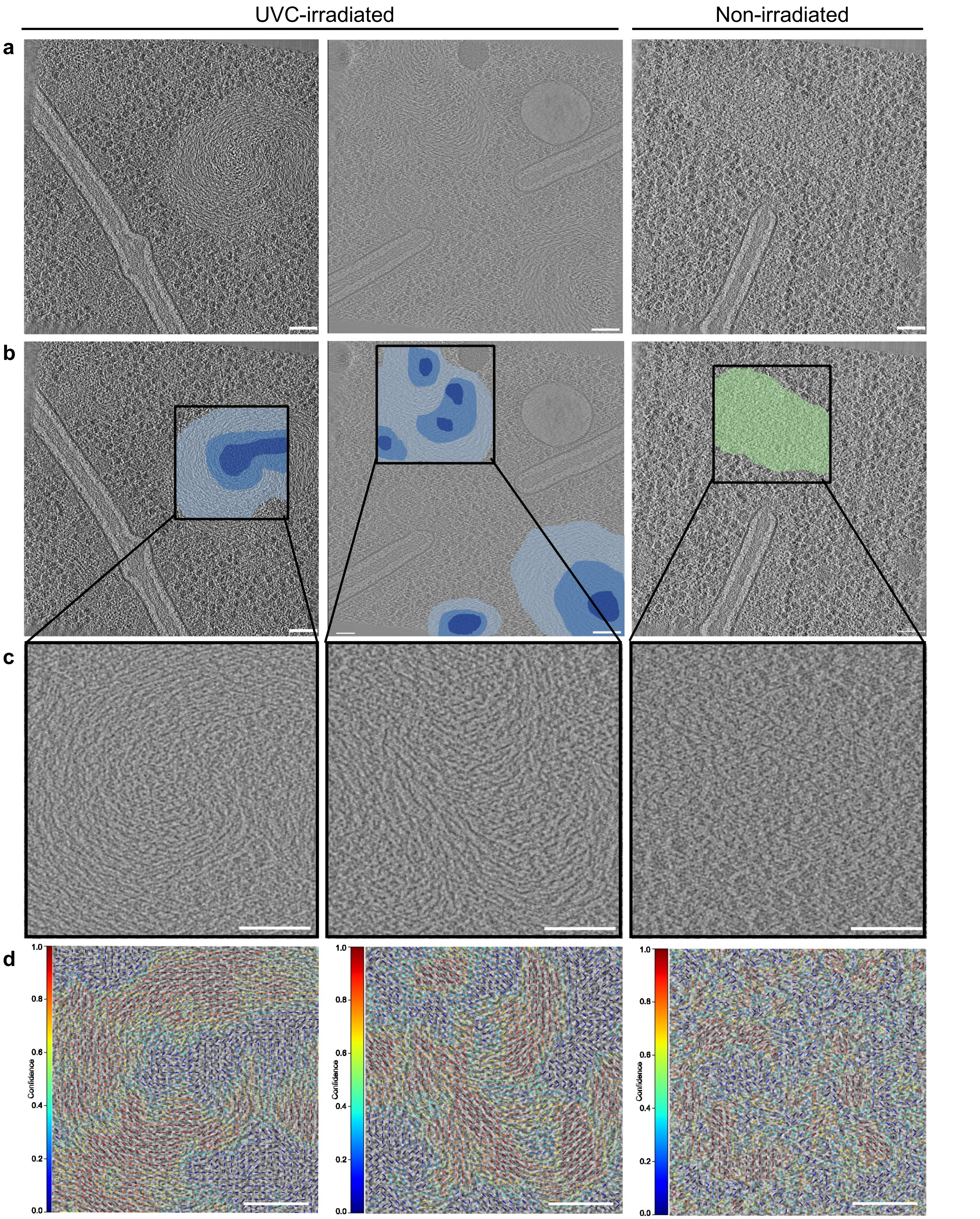


**Figure S10:** Analysis of the internal organisation and DNA pattern of UV-C irradiated (left and center) and non-irradiated (right) *D. radiodurans* nucleoids. (**a-b**). Slices through representative tomograms illustrating different nucleoid organisations without **(a)** and with **(b)** colouring of the nucleoid as a function of its DNA pattern (shades of blue for a double twist arrangement, and pale green for a loose DNA mesh). **(c-d)** Close-up views of the boxed areas in **b. (d)** Close-up views overlaid with the confidence scores of the corresponding orientation field, which exhibits a clear spiral arrangement in irradiated samples (left and center). In regions where DNA filaments are readily distinguishable and locally aligned, filament orientation can be reliably estimated, resulting in high confidence scores (red). Conversely, where filaments are less distinct or less ordered, orientation estimates are less reliable and confidence scores decrease (green/blue). Scale bars: 75 nm.
